# Task-Parametrized Dynamics: Representation of Time and Decisions in Recurrent Neural Networks

**DOI:** 10.1101/2025.09.15.676356

**Authors:** Cecilia Jarne, Ryeongkyung Yoon, Tahra Eissa, Zachary P. Kilpatrick, Krešimir Josić

## Abstract

How do recurrent neural networks (RNNs) internally represent elapsed time to initiate responses after learned delays? To address this question, we trained RNNs on delayed decision-making tasks with progressively increasing temporal demands, including binary decisions, context-dependent decisions, and perceptual integration. We analyzed trained networks using connectivity statistics, eigenvalue spectra, readout alignment, and low-dimensional population trajectories. Across tasks, networks converged to qualitatively distinct but behaviourally comparable dynamical solutions, including oscillatory and non-oscillatory (ramping/decaying) regimes, consistent with solution degeneracy. Population activity was well approximated by a low-dimensional subspace and distributed across recurrent units rather than localized to individual neurons. Readout alignment was strongly epoch-dependent: as required by the near-zero target output during that epoch, activity evolved largely in the readout-null subspace prior to response generation, and became increasingly aligned with the output dimension near decision time. In sign-symmetric tasks, trained networks preserved exact sign-flip equivariance inherited from architecture and training symmetry. Together, these results show that temporal and decision-related computations can emerge through multiple dynamical regimes, while maintaining structured low-dimensional representations and comparable behavioural performance, mirroring biological principles of degeneracy and functional redundancy.

## 1 Introduction

Timing is critical for humans and animals to navigate their environment and make the rapid choices necessary for survival (Paton and Buonomano 2018; Jazayeri and Shadlen 2010). Recurrent Neural Networks (RNNs) have become a central modelling framework in computational neuroscience, and have been used to uncover potential mechanisms that biological networks use to process temporal and sensory information and generate decisions (Mante et al. 2013; Russo et al. 2020; Laje and Buonomano 2013; Buonomano and Maass 2009). The ability of RNNs to generate rich, high-dimensional dynamics makes them particularly well-suited for understanding how neural networks can integrate multiple time-varying inputs and uncovering latent neural computations (Barak 2017; Maheswaranathan et al. 2019; Mastrogiuseppe and Ostojic 2018). For example, previous work has focused on how context and decision variables are represented in RNN population activity (Mante et al. 2013; Remington et al. 2018). Yet, there is still no consensus on how networks concurrently represent and update multiple dynamically evolving, task-dependent variables. Despite the possibility of high-dimensional RNN dynamics, many studies have shown that, after training, network activity converges to a small set of dominant trajectories or a low-dimensional manifold (Vyas et al. 2020). In particular, how RNNs can simultaneously encode elapsed time and decision variables during delayed decision-making tasks, and what underlying dynamical mechanisms allow them to do so, remains unclear.

How time is represented in RNNs and biological networks is not fully understood (Merchant et al. 2013a), since the encoding of time depends on both the specific cognitive process being modelled and how the task is parameterized. For instance, in RNNs trained to generate variable time intervals, temporal scaling (i.e., temporal compression/expansion of neural trajectories while preserving their shape) can depend on the strength of external input (Hardy et al. 2018). These findings are consistent with experiments where non-human primates trained to report time intervals show heterogeneous temporal dynamics within cortical populations but unified population trajectories that show comparable temporal scaling (Wang et al. 2018). This same population-level temporal scaling has also been observed in other brain regions, including the striatum (Mello et al. 2015), supplementary motor area (Merchant et al. 2013b) and hippocampus (MacDonald et al. 2011; Wang et al. 2018; Merchant et al. 2013b). Additionally, (Harvey et al. 2012) identified sequence generation as a timing mechanism in the cortex. Likewise, both RNNs and biological neural circuits can perform timing and decision-making tasks by engaging diverse computational mechanisms and neural population dynamics (Vyas et al. 2020), including ramping activity (Merchant et al. 2013b; Mendoza et al. 2018; Merchant and de Lafuente 2024), integration, oscillations, and sequence generation (Sussillo and Abbott 2009; Barak et al. 2013; Buzsáki and Wang 2012). Characterizing how different regimes of network activity and neural architectures emerge under different task conditions is essential for uncovering constraints on computational implementations in neural systems.

It is unclear whether neural networks converge to a single temporal representation or whether the encoding of time changes with task demands. We expect that the emergence of diverse temporal coding strategies is intimately tied to task complexity, which shapes the neural solution space in non-trivial ways. Increasing task complexity can either constrain the solution space, limiting the diversity of network dynamics that emerge during training (Huang et al. 2024), or result in a rougher optimization landscape with more local minima, potentially promoting a diversity in the dynamics of trained networks (Choromanska et al. 2014; Lyle et al. 2022; Nanda et al. 2023). Thus, it is necessary to determine which of these effects dominates under realistic training conditions.

Here, we ask how RNNs encode time when trained to perform decision-making tasks with varying demands. We begin with a simple binary decision-making task and progressively introduce additional complexity. This systematic increase in task complexity allows us to isolate how networks represent elapsed time and use these representations across tasks with varying demands. In particular, we ask how task-dependent delay periods (e.g., requiring a response after a delay in the absence of an external *cue*) are represented by population activity, how network activity concurrently reflects decision-specific computations, and, ultimately, how internal representations of decision variables are translated into responses. In short, how does a network represent time while simultaneously preparing and then executing a decision?

Previous work has systematically varied network and task parameters and showed that delay structures (fixed versus variable) and architectural choices jointly shape whether memory is implemented via sequences or persistent states (Orhan and Ma 2019). Here we take a complementary angle: rather than asking what form of short-term memory emerges, we ask how elapsed time itself is represented while the network concurrently encodes decision variables. By parameterizing families of delayed decision tasks (fixed vs. variable delays; amplitude- or interval-coded timing; integration), we show multiple recurrent networks (or replicas) converged to oscillatory, non-oscillatory (ramping/decaying) or mixed solutions with comparable behavioural performance. Crucially, mirrored responses to positive/negative stimuli were not imposed: with zero-mean initialization and an odd nonlinearity (*σ* = tanh()), the untrained network is sign-flip equivariant by construction; because biases remain fixed at zero throughout training, this architectural equivariance is preserved exactly. Consistent with this, responses were antisymmetric at the population level, and decision-related activity occupied low-dimensional subspaces. Thus, distinct dynamical regimes can implement the same computation, while population codes remain low-dimensional. Throughout this work, we use the term solution degeneracy to denote the existence of multiple internal dynamical implementations that achieve comparable behavioural performance despite differing recurrent dynamics and connectivity.

## 2 Methods

### 2.1 Model

We trained fully connected RNNs described by Eq. 1 on a sequence of tasks. The dynamics of the RNN model of *N* units is described in terms of the activity column vector **h**. We denote the recurrent weight matrix as **W**^Rec^, the input weight matrix with **W**^in^, and the bias term with a column vector **b**. The readout, **z**(*t*), of the network dynamics is given by Eq. 2, where **W**^out^ is the output weight matrix.

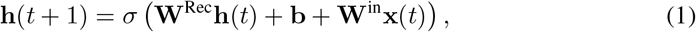

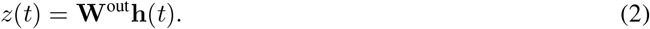

For our single-output readout, *z*(*t*) is scalar and this term enforces **z**(*t*) ≈ 0 prior to the response; equivalently, the hidden trajectories evolve roughly in the null space of the readout (orthogonal to **W**^out^) during this interval. We used a hyperbolic tangent for the activation function, *σ*. Panels (a) and (b) of Fig. 1 illustrate a representation of the network. Each network includes one or more input units, depending on the task, and a single output that presents the decision result based on the computations performed. Our study focused on relatively small networks, containing 100 units, with the weights represented in the connectivity matrices shown in Fig. 1b.

**Fig. 1.**
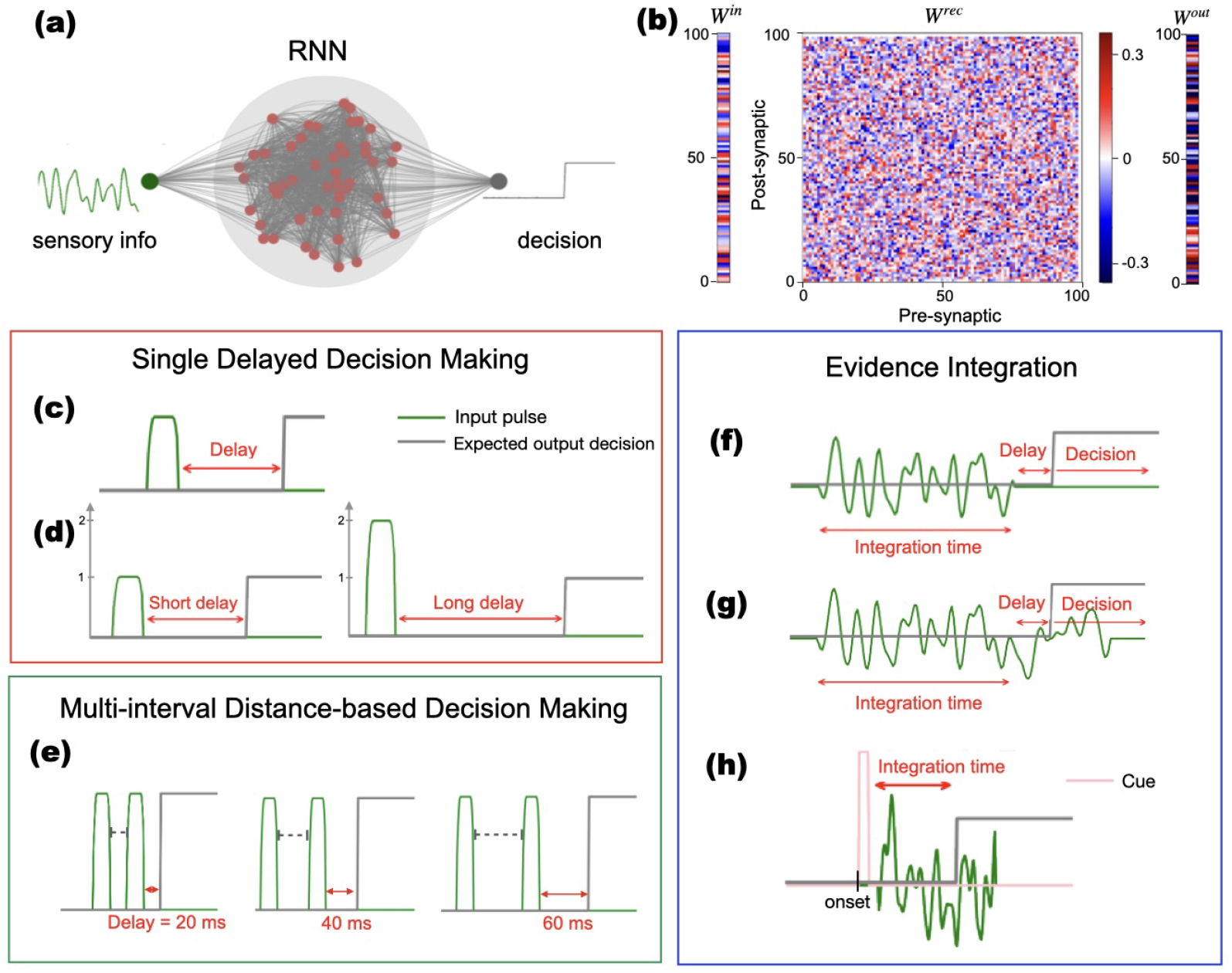
Task parametrization and network architecture. **(a)** Schematic representation of the RNN architecture, illustrating both the input and output signals. **(b)** Example of post-training RNN with input **W**^in^, recurrent, **W**^Rec^, and output, **W**^out^, layers. **(c)-(h)** Tasks schematics: **(c)** Simple Delayed Binary Decisions: Rectangular input pulses whose sign corresponds to an output decision. **(d)** Context-dependent Binary Decisions: Amplitude-modulated stimuli encoding short/long intervals. **(e)** Interval-based Context-dependent Decisions: temporal intervals are encoded via pulse separation. **(f)** Windowed Evidence Integration: Signal integration over a predefined window of evidence. **(g)** Continuous Evidence Integration: signal integration in a predefined window of a continuous evidence stream **(h)** Cued Evidence Integration: Amplitude-based cue defines the duration of evidence integration.

### 2.2 Training

For each of the tasks presented in the bottom panels of Fig. 1 and Table 1, we trained a set of 10 RNNs for each initial condition. To ensure our findings were robust across random initializations, we computed each statistic across the ensemble of trained RNNs and monitored changes in the metrics as new networks were added. We found that all relevant measures varied by less than 5% with the addition of each new network, indicating convergence. This approach follows a practice similar to that of (Pandarinath et al. 2018), who assessed the consistency of extracted dynamics across multiple model instantiations. During the training process, we allowed all input, recurrent, and output weights to change. Symmetry in the population responses was built in from the initialization (orthogonal or zero-mean random weights) and preserved during training. We employed a supervised learning method based on gradient descent using the adaptive minimization method Adam (Kingma and Ba 2014) with a batch technique. This method and variants have been successfully used to train RNNs on many tasks previously (Yang et al. 2019; Russo et al. 2018; Jarne and Caruso 2023; Jarne and Laje 2023).

**Table 1.**
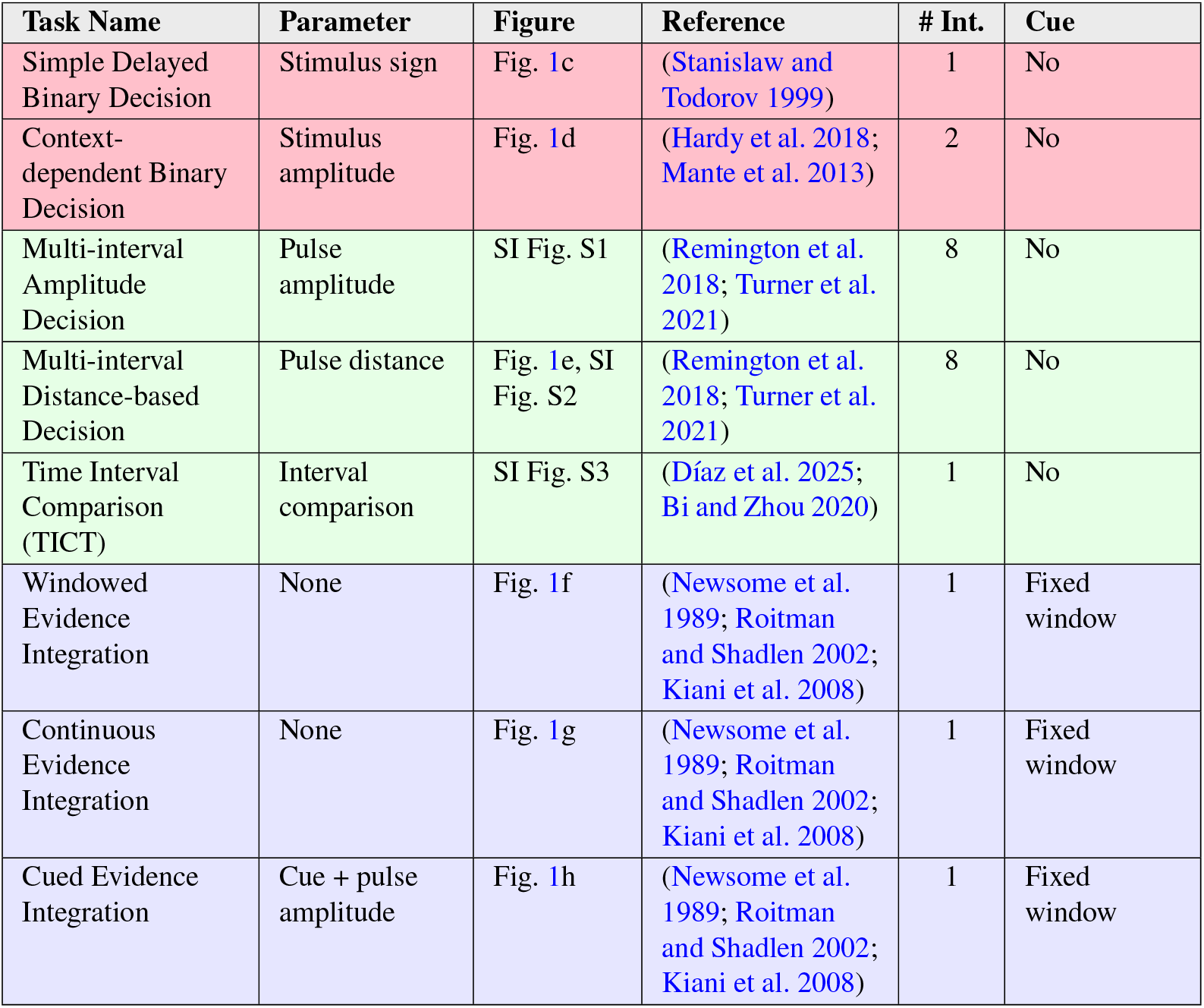
Summary of all tasks included in the study: parameters, figure for task representation, reference, number of intervals (# Int), and cue presence. Category 1 (red): simple decisions with single delays. Category 2 (green): binary decisions with multiple intervals. Category 3 (blue): perceptual integration tasks.

| Task Name | Parameter | Figure | Reference | # Int. | Cue |
| --- | --- | --- | --- | --- | --- |
| Simple Delayed Binary Decision | Stimulus sign | Fig. 1c | (Stanislaw and Todorov 1999) | 1 | No |
| Context-dependent Binary Decision | Stimulus amplitude | Fig. 1d | (Hardy et al. 2018; Mante et al. 2013) | 2 | No |
| Multi-interval Amplitude Decision | Pulse amplitude | SI Fig. S1 | (Remington et al. 2018; Turner et al. 2021) | 8 | No |
| Multi-interval Distance-based Decision | Pulse distance | Fig. 1e, SI Fig. S2 | (Remington et al. 2018; Turner et al. 2021) | 8 | No |
| Time Interval Comparison (TICT) | Interval comparison | SI Fig. S3 | (Díaz et al. 2025; Bi and Zhou 2020) | 1 | No |
| Windowed Evidence Integration | None | Fig. 1f | (Newsome et al. 1989; Roitman and Shadlen 2002; Kiani et al. 2008) | 1 | Fixed window |
| Continuous Evidence Integration | None | Fig. 1g | (Newsome et al. 1989; Roitman and Shadlen 2002; Kiani et al. 2008) | 1 | Fixed window |
| Cued Evidence Integration | Cue + pulse amplitude | Fig. 1h | (Newsome et al. 1989; Roitman and Shadlen 2002; Kiani et al. 2008) | 1 | Fixed window |

For tasks shown in Fig. 1c-e, the stimuli consisted of rectangular pulses with added random Gaussian noise amounting to 10% of the pulse amplitude or, if no stimulus was presented, only random low-amplitude noise (*<* 10% of stimulus amplitude). Such noise was added to generate trained RNNs that are robust to small fluctuations in input. For integration tasks, shown in Fig. 1f-h, the network was trained to integrate random signals during a time interval and report whether the integrated signal is positive or negative. The integration window can be defined by the time at which the input stops (f), a learned and internally represented duration (g), or cued by an amplitude-based secondary input (h). Within each trial, the stimulus onset time was randomized: The simulation started at time *t* = 0, and the stimulus or cue appeared at a randomly chosen time thereafter up to a maximum time *t* = *t*_*i*_. The target output completed the training set and varied depending on the task. The target was determined by the correct response given the integration of the input stimulus and the time at which a response needed to be generated. All tasks could be learned using 100 units and between 20 and 40 epochs over the training dataset (depending on the task), which contained 15,000 time series (trials), for example, for a simple Delayed Binary Decision task. A single epoch corresponds to one full pass through these 15,000 training trials.. Time series length varied between 350 and 500 time points, depending on the task (Supplementary Table S1).

The network was trained using the mean squared error (MSE) between the predicted output and the target signal, averaged across all time steps and across all trials within each training batch. For a batch of size *B* and a sequence length *T*, the loss is:

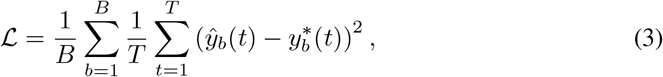

where (*B*) is the batch size and (*T*) the sequence length. *ŷ*(*t*) is the network output at time *t*, and 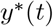 is the corresponding target. The target is set to 0 during the pre-decision period (*t* ≤ *T*_pre_) and to the required decision value (± 1 or a continuous value, depending on the task) during the post-decision period (*t* > *T*_pre_). No additional temporal weighting or separate normalization of the pre- and post-decision phases was applied; all time steps contributed equally to the loss. More details are shown in SI Table S1.

### 2.3 RNN initialization

We used two standard initialization schemes for the recurrent weight matrix, random normal and random orthogonal, to assess the sensitivity of task performance to initial conditions. For each scheme, we trained 10 RNN replicas on the same binary decision-making task and each of the tasks in Table 1. Despite differences in their learned connectivity matrices and eigenvalue spectra, all replicas achieved broadly similar performance levels, suggesting convergence in functional output, if not in internal representations (Jarne and Caruso 2023; Jarne and Laje 2023). Recurrent weights were initialized either from a zero-mean Gaussian distribution *N* (0, *σ*^2^) with 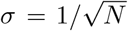, or from a random orthogonal matrix scaled by a gain parameter *g* = 1, resulting in an initial spectral radius close to unity. We considered zero biases, and used an odd nonlinearity (tanh()). This combination implies sign-flip equivariance at initialization: sign-reversed stimuli evoke mirror-reversed population responses. In bias-free networks, sign-flip equivariance is an architectural property of the deterministic dynamics. For matched sign-reversed inputs, the hidden trajectories satisfy *h*_−_(*t*) = − *h*_+_(*t*). For matched sign-reversed trial pairs, the identical (mirrored) noise realization was applied to both members of the pair, so that *h*_−_(*t*) = − *h*_+_(*t*) remain exact mirror images throughout the trial Because the training set and loss were sign-symmetric and we imposed no explicit symmetry constraint, gradient-based training preserved this architectural symmetry. With zero biases, an odd nonlinearity, and *h*(0) = 0, this equivariance is exact for any recurrent, input, and output weights: if *h*(*t*) solves Eq. 1 for input *x*(*t*), then − *h*(*t*) solves Eq. 1 for − *x*(*t*), since *tanh*() is odd and the recurrence contains no constant term, and by Eq. 2 the readout satisfies *z*(− *x*) = − *z*(*x*). Genuine breaking of this equivariance therefore requires an even component in the dynamics, such as a nonzero bias, a non-odd activation function, an initial state that is not itself sign-reversed, or independently (non-mirrored) sampled noise between paired trials. In our simulations, matched +*x/* − *x* trial pairs received identical (mirrored) noise realizations, so no such source of asymmetry was present, and the equivariance holds exactly for every trial pair.

The expected sign-flip equivariance was maintained exactly throughout training, to floating-point precision. The symmetry analysis therefore serves as a validation that optimization preserves the architectural constraints imposed by zero biases, odd activation functions, and sign-symmetric training data, rather than as evidence for an additional computational principle.

### 2.4 Task parametrization

We considered several versions of delayed binary response tasks with a focus on temporal representation in the networks. The expected responses and the length of the response delay differed depending on the specific task and parameters, as explained in the next sections and summarized in Table 1. The different response delay times corresponded to the elapsed time that had to be learned by the network.

The tasks can be grouped into three categories, which are colour-coded in Table 1. Category 1 comprises tasks that require a response after a learned delay. Category 2 comprises tasks that involve a binary decision that needs to be reported after one of multiple learned time intervals. Category 3 comprises tasks that require the integration of perceptual information over a learned time period, followed by a response after a learned, but unsignalled, time interval. Target intervals were drawn from a discrete set *T*_*i*_ ∈{*T*_1_, …, *T*_8_}, uniformly spaced 20 to 160 in steps of 20. In amplitude-cued tasks, each *T*_*i*_ is associated with a stimulus amplitude *A*_*i*_, defining a one-to-one mapping *T*_*i*_ = *f* (*A*_*i*_), from 25 to 200 steps. In pulse-cued tasks, *T*_*i*_ is directly specified by the temporal separation between consecutive input pulses. Amplitudes *A*_*i*_ were linearly spaced and in the range [1, 8].

#### Simple Delayed Binary Decision Making

We first considered a task where the network learns to report the sign of an input signal after a single, learned time interval. This simple delayed report task is based on the two-alternative forced-choice (2AFC) paradigm, which requires selecting between two predefined options in response to a stimulus (Stanislaw and Todorov 1999). We used a single square pulse as input, with the sign of the pulse determining the sign of the correct response. Pulse amplitude was held fixed, but was irrelevant for the task. The network was trained to provide a constant output which matches the sign of the input signal after a fixed, learned delay (see Fig. 1c). The delay was learned, and no cues were provided to signal the report time.

#### Context-dependent Binary Decision Making

The context-dependent binary decision-making task reparameterizes the simple delayed binary decision-making task to include two different delay intervals that are signalled by the stimulus amplitude (see Fig. 1d). When the signal amplitude is low (high), the delay time is short (long) and equals 50 (100) time units. This design allowed us to ask how the sign of the input signal and elapsed time were represented in the activity of the trained network.

#### Multi-interval Amplitude-based Decision Making

For the multi-interval amplitude-based decision-making task, we used eight distinct stimulus amplitudes, each corresponding to a different time at which a response was to be provided (SI Fig. S1). The target output remained binary (positive or negative, depending on the input signal’s sign), but the network had to interpret the stimulus amplitude and track elapsed time to provide a response at the correct, unsignalled time. Each interval *T*_*i*_ defines the target delay between stimulus onset and required response time. This design allowed us to ask how the trained network represents the correct response and translates it into a response after a signalled time interval.

#### Multi-interval Distance-based Decision Making

In this task, the required response time was encoded by the time between two consecutive input pulses (See Fig. 1e) and (SI Fig. S2). Unlike amplitude-based encoding, this approach establishes a more explicit mapping between the input interval and the output timing. To compare these two representations of time, we trained recurrent neural networks on both tasks using the same set of eight distinct intervals. This allowed us to ask how networks represent elapsed time when the length of a time interval is represented in a different way by the input. This task is conceptually related to the Ready-Set-Go paradigm (Remington et al. 2018; Turner et al. 2021), in that the inter-pulse interval *T* sets the delay before a response is required. Unlike Ready-Set-Go, however, the network is not required to reproduce *T* in its output: the required response is a constant ± 1 decision whose sign reports the stimulus polarity, while *T* only determines the timing at which that decision must be reported. We use eight distinct values of *T* to contrast this distance-based timing cue with the amplitude-based timing cue described above.

#### Time Interval Comparison Task (TICT)

Here, the network compared two temporal intervals (int1 vs. int2) which were presented sequentially and separated by a fixed delay. The target response indicated whether the first or second interval was longer, making this a version of a standard interval discrimination task (Díaz et al. 2025; Bi and Zhou 2020), (see SI Fig. S3).

#### Windowed Evidence Integration (Perceptual Decision Making)

Here, the network was required to integrate an input signal over a learned time window. The sign of this integral agreed with the sign of the required response, and the response needed to be reported after a fixed time interval (Fig. 1f). No explicit cue was provided to signal when integration needed to cease and a response needed to be generated, but the response needed to be generated at a fixed window after the cessation of the input.

#### Continuous Evidence Integration

This task was similar to the Windowed Evidence Integration task, but the input signal does not end before a response is required (Fig. 1g). Rather, the network is trained to respond after a fixed amount of time. Thus, the network has to track the elapsed time along with the decision variable.

#### Cued Evidence Integration

This was an extension of the Continuous Evidence Integration task, with an additional cue at trial onset that signalled the duration of the integration time. If there was no cue signal, the network’s output remained zero (Fig. 1h). Thus, the network needed to represent elapsed time and translate the decision variable to a response after a signalled time interval that varied from trial to trial.

### 2.5 Analysis

Our analyses are summarized in Fig. 2. We first examined connectivity statistics, including weight distributions and their moments, to quantify training-induced changes (Fig. 2a). Second, we computed the eigenvalue spectra of the recurrent weight matrix **W**^Rec^ to characterize global dynamical regimes (Fig. 2b). Throughout this work, a dynamical regime refers to the dominant qualitative pattern of recurrent population activity expressed by a trained network during task execution. Although eigenvalue spectra do not capture input- or state-dependent dynamics in nonlinear RNNs (Schuessler et al. 2020; Krause et al. 2022; Valente et al. 2022), they provide informative summaries of how training reshapes connectivity structure (Rajan and Abbott 2006; Rivkind and Barak 2017). Networks with eigenvalues outside the unit circle with substantial imaginary components tended to support oscillatory dynamics, while those with a dominant real eigenvalue produce ramp-like or decaying activity (Sussillo and Barak 2013; Jarne 2022); such interpretations are supported by linearizations around fixed points (Sussillo and Barak 2013). Third, we analyzed temporal activity traces at both single-unit (Fig. 2c) and population levels across task conditions. Fourth, we investigated low-dimensional population trajectories (Fig. 2d) using principal component analysis (PCA), enabling visualization of task-dependent dynamics.

**Fig. 2.**
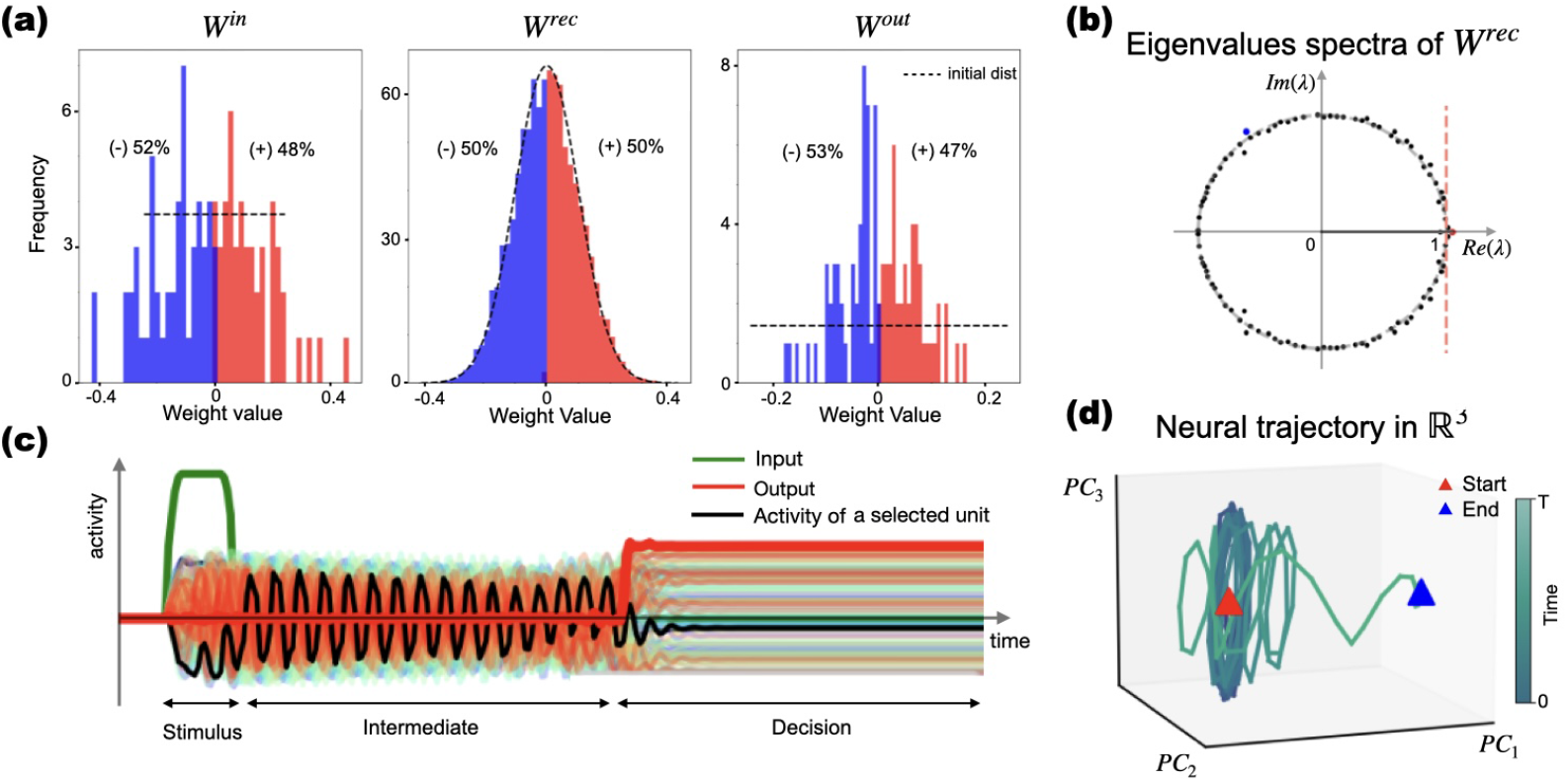
Framework for analyzing trained RNNs. Panels show an illustrative example from a single network trained on the Simple Delayed Binary Decision Making task (orthogonal initialization). **(a)** Histograms of weight distributions after training: Input **W**^in^, recurrent, **W**^Rec^, and output, **W**^out^. Histograms are compared to their respective initialization distributions (e.g., Gaussian or orthogonal). These examples illustrate how training modifies network connectivity. **(b)** Spectral analysis of network dynamics: Eigenvalues of the **W**^Rec^ matrix are shown in the complex plane. The spectrum relates to the stability and oscillatory activity supported by the network. **(c)** Temporal activity across task phases: Example activity traces from a single trial. The green line depicts the stimulus input, the red line shows the network’s output. Colored traces represent the activity of individual units during the stimulus, intermediate, and decision phases. The black line indicates the activity of one highlighted unit. **(d)** Low-dimensional representation of neural trajectories: Neural activity of the network during a trial was projected onto the first three principal components. The red triangle marks the start of the trajectory, and the blue triangle marks the end.

In addition, we performed ablation analyses to assess robustness. We ablated internal (recurrent) units by removing the unit from the recurrent graph – i.e., zeroing its incoming and outgoing entries in **W**^Rec^, zeroing its readout weight in **W**_out_, and clamping its state to 0 during simulations – and then examined the resulting changes in network activity and the spectrum of the modified recurrent connectivity.

Trained networks were also classified into dynamical regimes using a dual criterion that combines spectral and activity-based metrics. The spectral Oscillation Index OI = | Im(*λ*^∗^)| /| *λ*^∗^| was computed from the dominant eigenvalue of **W**^Rec^. The activity criterion was based on the peak-to-background ratio PBR of the first principal component during the delay epoch and a ramp index (absolute Pearson correlation with time). Full details of the classification procedure, including the thresholds, are provided in SI in the Supplementary Methods section.

#### 2.5.1 Generalized correlation and readout alignment

We also analyzed whether population activity was structured relative to the output readout. To capture interactions between units, we computed the correlation matrix of neural activity across time. We used the generalized correlation measure suggested by Schuessler et al. (2024), as a descriptive measure to characterize how the relationship between population activity and the readout evolves across task epochs within a given trained network. This metric provides a mathematically consistent measure of alignment between the output matrix weights (**W**^out^) and neuronal activity: Consider the neural activity of *N* neurons at *P* time points of the time series stacked into the vector **X**. Then, the corresponding *D*-dimensional output is summarized in the *D* × *P* matrix **Z**, from Eq. 2. The generalized correlation in Schuessler et al. (2024) is defined as:

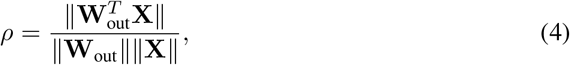

where ∥ · ∥ denotes the Frobenius norm. This quantity measures the global alignment and collective contribution of the neural population to the output. This allows us to classify the computational strategies employed by trained networks. We use this metric descriptively to track how alignment between population activity and the readout evolves across task epochs, rather than to classify networks as operating in aligned or oblique regimes. In our setting, the metric primarily captures the temporal recruitment of the readout dimension during task execution.

#### 2.5.2 Population geometry and principal angles

To characterize the geometry of population representations, we analyzed low-dimensional trajectories derived from trial-averaged hidden-state activity. For each trial, the activity was segmented into task-relevant stages—pre-stimulus, stimulus, intermediate, delay/decision, and full trial—based either on threshold detection of the input and output signals or on the task’s known deterministic structure. Fig. 2c shows an example of unit activity (black line) together with the input stimulus (green line) and network output (red line), highlighting the temporal boundaries of these stages.

From these stage-segmented trajectories, we then compared the population subspaces occupied under different task conditions (e.g., positive vs. negative response, different stimulus amplitudes or temporal intervals, long vs. short pulses). For each condition, principal component analysis (PCA) was applied to the time-by-neuron activity matrix within each stage, and the leading three principal components were retained, yielding a low-dimensional subspace that captures the dominant directions of population variance (Fig. 2d). The degree of alignment between the subspaces of different conditions was quantified by the principal angles, computed via singular value decomposition of the subspace bases. Larger angles indicate more orthogonal representations, while smaller angles suggest that the networks encode the conditions in similar population modes (Elsayed et al. 2016).

To obtain robust estimates, the same analysis was performed independently on multiple network replicas trained from different random initializations. For each condition pair and epoch, the mean principal angles and their standard deviations across replicas were computed. The explained variance ratio (EVR) of the leading PCs, assessed on the full trial activity, was also recorded and compared between conditions. Results are visualized, in SI, as bar plots showing the angle per PC, with overlaid jittered points representing individual replicas. Heatmaps summarized the mean angles across epochs and PCs. Separate EVR bar plots, together with numerical summary tables, allowed direct comparison of the variance captured by the dominant modes for each condition. This approach enabled a systematic assessment of how task variables (stimulus sign, magnitude, temporal structure, and pulse duration) modulate the geometry of the representational subspaces and their alignment, and whether the degree of orthogonality depends on the processing phase.

#### 2.5.3 Symmetry Index computation

To quantify the degree of symmetry, we defined a symmetry index (SymIdx). Let *x*^(+)^(*t*) ∈ ℝ^*N*^ and *x*^(−)^(*t*) ∈ ℝ^*N*^ denote the hidden-state trajectories for a matched +/ −trial pair. The instantaneous symmetry error is

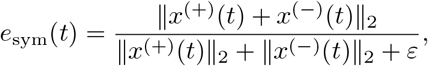

with *ε* > 0 for numerical stability. We summarize over time points *T* by

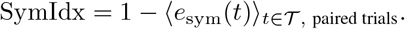

SymIdx = 1 indicates perfect sign-flip equivariance. Across all tasks and trained networks, SymIdx = 1 to floating-point precision, confirming exact sign-flip equivariance at the population level, as expected for a bias-free network with an odd nonlinearity and mirrored trial-pair noise. We emphasize that this analysis is intended as a validation of the expected architectural symmetry, rather than as a novel mechanistic finding. Its purpose is to verify that this equivariance, implied by the model assumptions, is preserved exactly by the training protocol used here

#### 2.5.4 Non-normality of networks

In addition to the spectral analyses above, we quantify non-normality using Henrici’s departure index (Eq. 5), which measures how far *W* ^Rec^ deviates from a normal matrix:

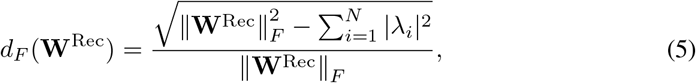

where ∥ · ∥_*F*_ denotes the Frobenius norm and *λ*_*i*_ are the eigenvalues of **W**^Rec^. This scalar metric globally quantifies how far **W**^Rec^ departs from a normal matrix, capturing both the eigenvalue distribution and the non-commutativity between **W**^Rec^ and its transpose.

A non-normal matrix (*d*_*F*_ > 0) is a necessary condition for transient amplification in linear systems, and has been shown to facilitate transient trajectories in recurrent networks (Bondanelli and Ostojic 2020). Thus, Henrici’s index complements eigenvalue spectra by indicating whether the connectivity structure *could potentially* support transient amplification. It has been successfully applied in recent studies of transient amplification in recurrent systems (e.g., Asllani et al. (2018)) and serves here as a concise, comparable summary statistic across networks.

We chose Henrici’s index because it is well-suited to our comparative goal: it yields a single interpretable scalar, is inexpensive to compute, and does not require choosing an epoch-specific operating point or computing Jacobians along trajectories. Alternative measures of non-normality have been proposed in the context of neural dynamics (e.g., (Bondanelli and Ostojic 2020)), but Henrici’s index offers a practical balance between informativeness and computational cost for our cross-network comparisons.

However, Henrici’s index is not a direct measure of transient amplification in the actual nonlinear, input-driven network. It does not capture input- or state-dependent effective dynamics, and a full characterization of transient amplification would require analysis of the state-dependent Jacobian (Schuessler et al. 2020; Krause et al. 2022). Therefore, we interpret Henrici’s index as a coarse global descriptor of connectivity structure, and we avoid strong mechanistic claims that would require a more detailed input-dependent analysis.

## 3 Results

We trained fully connected RNNs on a set of delayed decision-making tasks of increasing complexity (Table 1) and analyzed the resulting dynamics using eigenvalue spectra, single-unit activity traces, low-dimensional population trajectories, and readout-alignment metrics. Our central finding is that initialization scheme does not deterministically specify which dynamical regime, oscillatory or non-oscillatory (ramping/decaying), a network settles into, and behavioural performance is largely regime-invariant (Section 3.1). This degeneracy is not unconstrained: a logistic classifier combining spectral descriptors of the final recurrent connectivity with the initialization scheme discriminates oscillatory from other networks above a task-stratified permutation null (balanced accuracy 0.68, *p* = 0.005), even though no individual descriptor survives correction for multiple comparisons (Section 3.8).

Against this backdrop of regime variability, population-level symmetry behaves in the opposite way: zero-bias initialization with an odd nonlinearity (tanh()) and sign-symmetric training preserve exact sign-flip equivariance (SymIdx = 1 to floating-point precision) across all tasks and regimes, despite noisy inputs and stochastic optimization (Section 3.2). Symmetry is therefore a robust, regime-independent property of the learned solutions: it functions as an internal validation of architectural preservation rather than as a task-dependent discovery in its own right.

Finally, we characterized how population activity is temporally organized relative to the readout: alignment with **W**^out^ remains low throughout the stimulus and delay epochs and increases sharply only near the decision report time (Section 3.3). Because the target output is constrained to be near zero before the response period, this pattern follows directly from the task and loss design; we therefore treat it as a descriptive tool that, together with the non-normal, distributed recurrent reconfiguration reported in Sections 3.4–3.6, characterizes how the underlying degenerate dynamical solutions are translated into behavior.

### 3.1 Distinct Activity Patterns Result in Equivalent Task Performance

#### Distinct dynamical regimes emerge across networks trained on identical tasks

Trained RNNs converged to qualitatively distinct temporal dynamics while achieving comparable behavioural performance on the same delayed decision-making tasks. Some networks exhibited oscillatory activity during the delay period, whereas others displayed predominantly non-oscillatory ramping or decaying transients; a subset exhibited mixed dynamical signatures (Table 2). These regimes were reflected in the eigenvalue spectra of the recurrent connectivity matrix (Fig. 3a), in the temporal activity patterns of individual units despite similar input-output behaviour (Fig. 3b), and in the geometry of the population trajectories projected onto the first three principal components (Fig. 3c). Across regimes, activity converged toward stable fixed points after the response epoch.

**Table 2.** Classification of trained networks by dynamical regime and initialization scheme together with task performance metrics ((A) and (B)). For each task and initialization condition, 10 networks were trained independently. Networks were classified as oscillatory, ramping, or mixed according to dominant activity and spectral signatures. Reported values correspond to mean ± standard deviation across trained networks. OI: Oscillation Index;logPBR: log_10_(PBR); RampIdx: ramping activity index.

| (A) Dynamical Regime Classification |  |  |  |  |  |  |  |
| --- | --- | --- | --- | --- | --- | --- | --- |
| Task | Init. | Osc. | Ramp | Mixed | OI | logPBR | RampIdx |
| Simple Delayed Binary DM | Normal | 8/10 | 1/10 | 1/10 | $0.52 \pm 0.28$ | $2.47 \pm 0.89$ | $0.54 \pm 0.32$ |
| | Orthogonal | 9/10 | 0/10 | 1/10 | $0.60 \pm 0.32$ | $1.73 \pm 0.35$ | $0.44 \pm 0.25$ |
| Context-dependent Binary DM | Normal | 5/10 | 1/10 | 4/10 | $0.27 \pm 0.34$ | $2.41 \pm 1.00$ | $0.47 \pm 0.29$ |
| | Orthogonal | 6/10 | 1/10 | 3/10 | $0.61 \pm 0.33$ | $1.42 \pm 0.54$ | $0.43 \pm 0.29$ |
| Multi-interval Amplitude-based | Normal | 5/10 | 0/10 | 5/10 | $0.42 \pm 0.39$ | $2.29 \pm 0.80$ | $0.85 \pm 0.18$ |
| | Orthogonal | 6/10 | 2/10 | 2/10 | $0.52 \pm 0.42$ | $1.27 \pm 0.44$ | $0.90 \pm 0.14$ |
| Multi-interval Distance-based | Normal | 5/10 | 3/10 | 2/10 | $0.43 \pm 0.34$ | $1.62 \pm 0.82$ | $0.95 \pm 0.04$ |
| | Orthogonal | 6/10 | 2/10 | 2/10 | $0.47 \pm 0.38$ | $1.34 \pm 0.43$ | $0.91 \pm 0.09$ |
| Windowed Evidence Integration | Normal | 9/10 | 0/10 | 1/10 | $0.65 \pm 0.30$ | $2.78 \pm 0.46$ | $0.95 \pm 0.01$ |
| | Orthogonal | 6/10 | 0/10 | 4/10 | $0.42 \pm 0.38$ | $2.43 \pm 0.51$ | $0.95 \pm 0.01$ |

**Table 2:** Classification of trained networks by dynamical regime and initialization scheme together with task performance metrics ((A) and (B)).
| (B) Task Performance |  |  |  |  |
| --- | --- | --- | --- | --- |
| Task | Init. | Accuracy | Timing Error | MSE |
| Simple Delayed Binary DM | Normal | $1.000 \pm 0.000$ | $0.15 \pm 0.20$ | $0.001 \pm 0.001$ |
| | Orthogonal | $1.000 \pm 0.000$ | $0.00 \pm 0.01$ | $0.000 \pm 0.000$ |
| Context-dependent Binary DM | Normal | $1.000 \pm 0.000$ | $2.60 \pm 2.26$ | $0.009 \pm 0.009$ |
| | Orthogonal | $1.000 \pm 0.000$ | $0.20 \pm 0.22$ | $0.001 \pm 0.001$ |
| Multi-interval Amplitude-based | Normal | $0.999 \pm 0.002$ | $8.01 \pm 8.71$ | $0.018 \pm 0.017$ |
| | Orthogonal | $0.996 \pm 0.013$ | $5.86 \pm 4.28$ | $0.007 \pm 0.002$ |
| Multi-interval Distance-based | Normal | $1.000 \pm 0.000$ | $2.10 \pm 0.95$ | $0.004 \pm 0.001$ |
| | Orthogonal | $0.965 \pm 0.105$ | $5.27 \pm 14.26$ | $0.003 \pm 0.006$ |
| Windowed Evidence Integration | Normal | $0.932 \pm 0.038$ | $4.30 \pm 1.54$ | $0.071 \pm 0.007$ |
| | Orthogonal | $0.970 \pm 0.007$ | $4.34 \pm 1.41$ | $0.068 \pm 0.005$ |

**Fig. 3.**
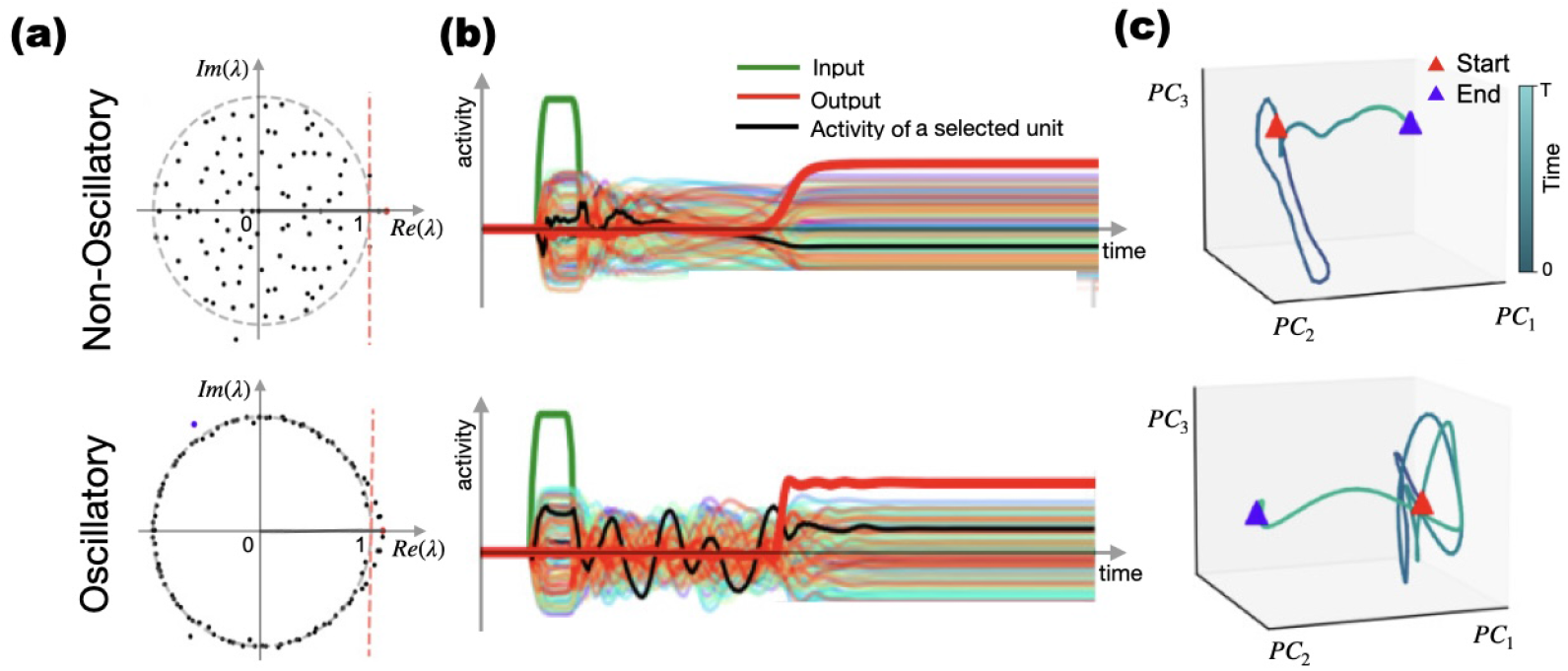
Representative examples of oscillatory and non-oscillatory dynamical regimes emerging from different initialization schemes. **(a)** Eigenvalue spectra of two RNNs showing non-oscillatory (top) and oscillatory (bottom) dynamics. **(b)** Input (green), output (red), a selected unit activity (black) and unit activity (colored) traces from each network. **(c)** Low-dimensional projection of the neural activity (top 3 PCs) showing convergence toward the response fixed point after a non-oscillatory transient (Top) and multiple oscillations (Bottom). Red: start, purple: end.

Networks initialized with random normal and orthogonal weight matrices could converge to different dynamical regimes despite identical task demands. Representative examples are shown in Fig. 3

Similar dynamical diversity was observed across more complex tasks. In the Multi-interval Amplitude-based Decision Making task, oscillatory solutions represented elapsed time through the number of oscillation cycles, while trajectories formed clustered structures in phase space associated with different delay intervals (SI Fig. S1a-b). Likewise, SI Fig. S2 shows two networks trained on the same Multi-interval Decision Making task that converged to different temporal strategies: one displaying predominantly ramping dynamics and the other oscillatory dynamics. These distinct regimes produced qualitatively different trajectories in the low-dimensional PCA subspace (SI Fig. S2c). Similar organizational principles extended to contextual integration tasks, where cue and non-cue conditions occupied partially orthogonal subspaces in PCA space (SI Fig. S4).

Across tasks, trained RNNs converged to oscillatory, non-oscillatory, or mixed dynamical regimes with comparable behavioural performance. These findings indicate that relatively simple RNNs can implement time-keeping through multiple qualitatively distinct yet functionally equivalent dynamical solutions, consistent with temporal coding motifs reported in biological neural systems.

To quantify these regimes systematically, we classified each trained network using a dual activity-spectral criterion (Table 2). The spectral criterion computes the Oscillation Index OI = |Im(*λ*^∗^)| /| *λ*^∗^|, where *λ*^∗^ is the dominant eigenvalue (largest modulus) of the recurrent weight matrix **W**^Rec^. Activity-based classification used a fixed post-pulse window starting immediately after the input pulse offset and during the delay epoch. On each trial, we injected a smoothed square pulse (Hann-windowed, ± 1 amplitude, with 5% additive Gaussian noise) and recorded the hidden-state trajectories. To avoid cancellation by sign-flip symmetry, only trials with positive-polarity input were averaged. From the averaged trajectory, the first principal component (PC1) of the delay-window activity was extracted, linearly detrended, and its power spectral density was estimated via Welch’s method. The peak-to-background ratio was computed as PBR = max_*f>*0.10 cyc/step_ *P* (*f*) / median_*f>*0.10 cyc/step_ *P* (*f*), and the log_10_ value was taken. A ramp index was defined as the absolute Pearson correlation between PC1 and time over the same window. We report log_10_ PBR. Networks were classified as **Oscillatory, Ramping/Decaying**, or **Mixed** based on both criteria. As complementary measures, we also computed a ramp index (absolute Pearson correlation between time and the raw PC1 projection) and the spectral OI.

The distribution of dynamical regimes varied across tasks and initialization schemes (Table 2). Orthogonal initializations increased the prevalence of oscillatory solutions in several tasks, although all initialization conditions remained capable of producing oscillatory, ramping/decaying, or mixed regimes with comparable behavioural performance.

### 3.2 Architectural Symmetry is Preserved During Training

Because we initialize all weights with normal or orthogonal distribution, set all biases to zero, and use the odd nonlinearity tanh(), the untrained network is sign-flip equivariant: a stimulus − *x* produces the negative of the activity evoked by +*x*. Training on symmetric binary tasks preserves exact sign-flip equivariance (SymIdx = 1 to floating-point precision). We observed mirror-symmetric unit activations for opposing stimuli across all tasks (Fig. 4, Fig. 5, SI Animations 1, 2, and 3), and opposite decisions traced mirrored trajectories in PCA space (Fig. 4b). This symmetry is a direct consequence of keeping biases fixed at zero. To test whether this symmetry depended specifically on the zero-bias constraint, we repeated training with trainable biases. In these networks, symmetry was partially but systematically broken SymIdx ≈ 0.77–0.80 (*e*_sym_ ≈ 0.20–0.23), confirming that trainable biases constitute the primary mechanism through which symmetry is broken in the present architecture. These observations confirm that the expected sign-flip equivariance imposed by the network architecture is preserved throughout training. The symmetry analysis therefore serves primarily as a validation that the learned population dynamics remain consistent with the architectural constraints, rather than as evidence for a distinct computational mechanism.

**Fig. 4.**
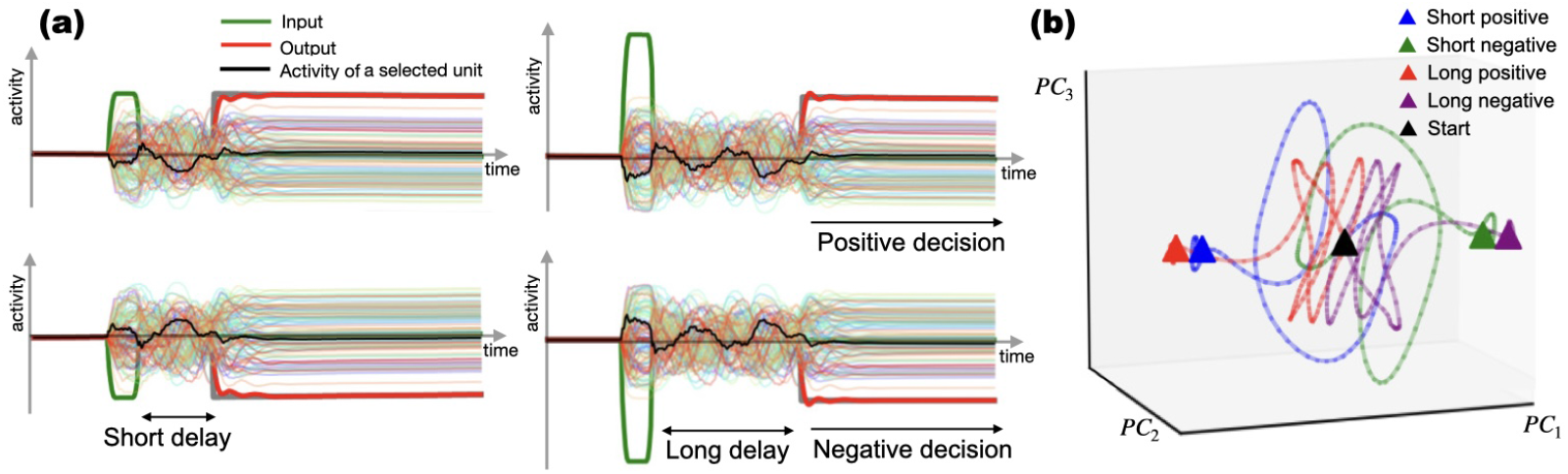
Symmetric internal dynamics in context-dependent binary decision making task **(a)** Neural responses to stimuli of equal amplitude and opposite sign produce mirror-symmetric unit activity across conditions. Example neural activity from multiple neurons is shown as thin traces. The black trace shows an example unit where the response inverts sign with stimulus polarity, demonstrating a lack of stimulus-specific specialization. **(b)** PCA trajectories for four conditions (“Short/Long” or “Positive/Negative”). Trajectories for opposite-sign stimuli converge to mirrored regions in state space, with consistent clustering across delay intervals.

**Fig. 5.**
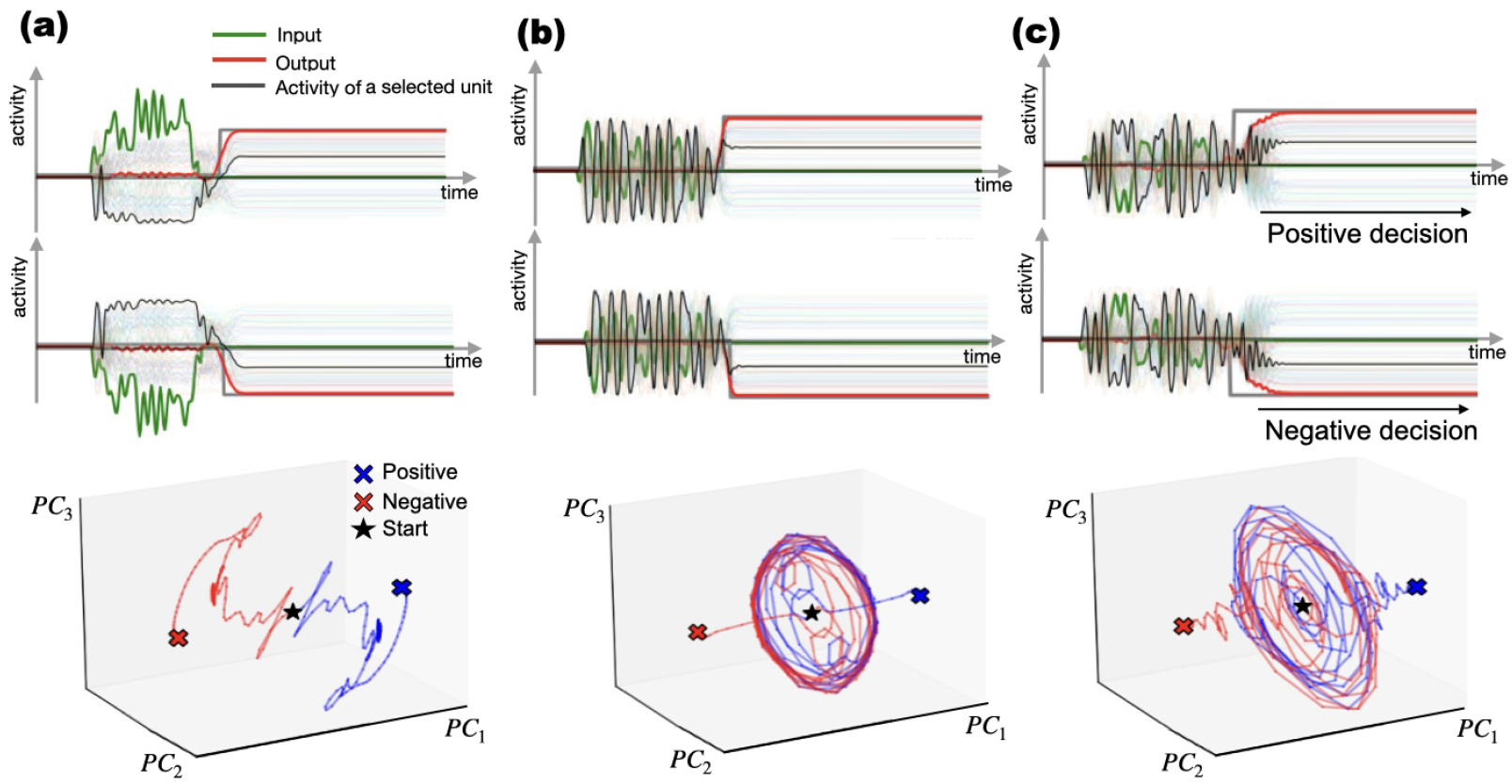
Internal dynamics during Windowed Evidence Integration task. Three different examples (columns **a, b** and **c**) of network activity in time (top, middle) and projected activity using PCA (bottom). The trajectories depend on the characteristics of the signal being integrated. When the random signals are equal and opposite, a symmetrical response is produced. This is also evident in the PCA graph, where the mirror-opposed trajectories indicate that the behaviour of all units reflects this pattern.

### 3.3 Temporal Decoupling Segregates Sensory Encoding and Decision Execution

To characterize how trained networks transition from sensory encoding to behavioural output, we quantified the temporal alignment of recurrent activity with the input and readout dimensions. Because the target output remains near zero before the decision epoch, strong pre-decision alignment with **W**^out^ is not expected from the task design itself. The relevant dynamical feature is therefore the sharp increase in readout alignment at decision time.

Input-output alignment, defined by Eq. 4, shifts dynamically across task phases (Fig. 6). For the simple decision-making task, during stimulus presentation, the units’ activity was strongly correlated with input weights,**W**^in^, (positive: 0.523 ± 0.091; negative: 0.515 ± 0.093), reflecting direct sensory encoding (Fig. 6b). It then decreases sharply during the intermediate (positive: 0.181 ± 0.016; negative: 0.180 ± 0.025) and decision periods (positive: 0.052 ± 0.002; negative: 0.055 ± 0.002). A similar temporal structure was observed in the integration task. Input alignment remained distributed across the signal and intermediate periods (positive: 0.292 ± 0.039 and 0.295 ± 0.021; negative: 0.291 ± 0.040 and 0.295 ± 0.029), but dropped during the decision period (positive: 0.051 ± 0.006; negative: 0.055 ± 0.005).

**Fig. 6.**
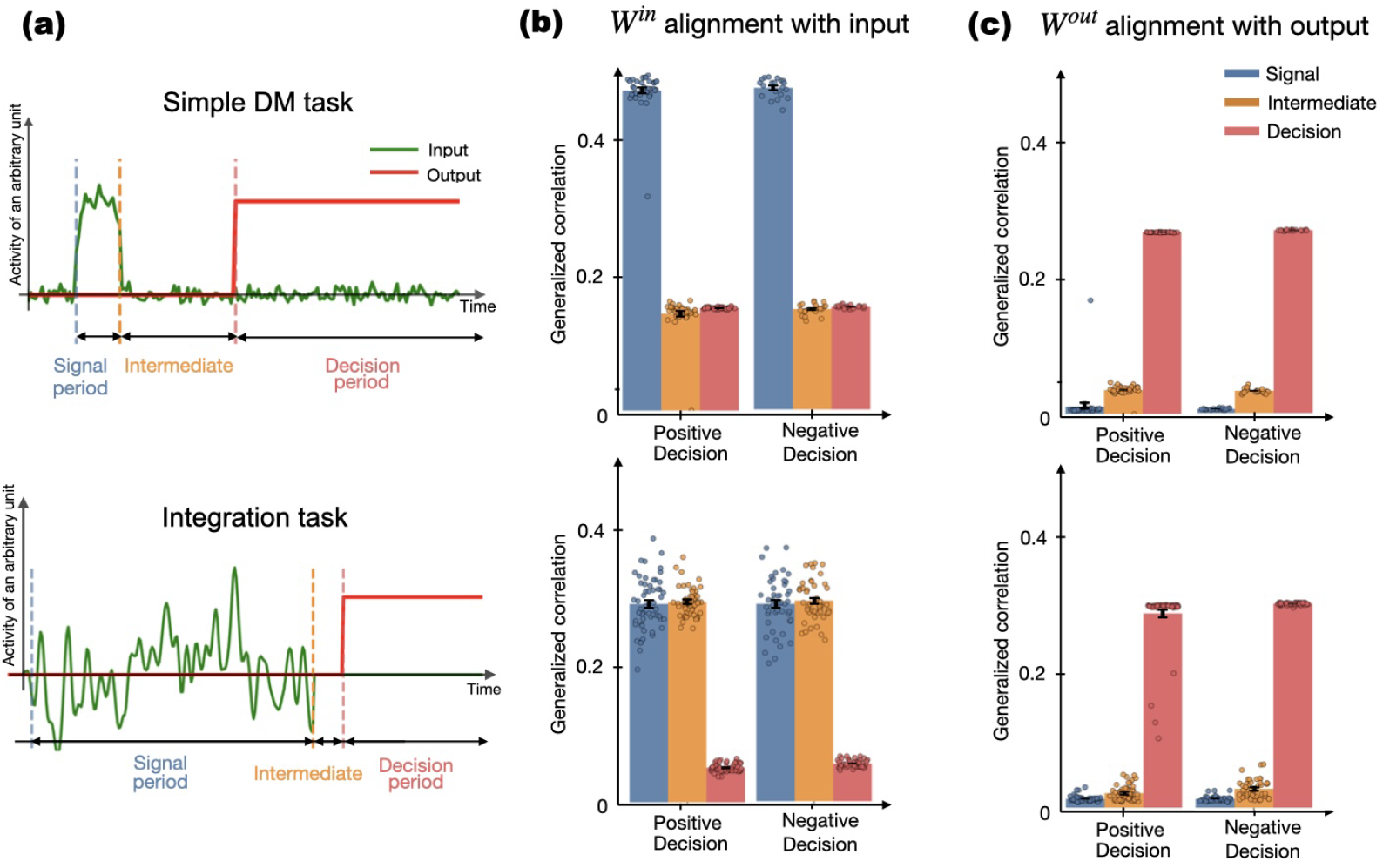
Input-output alignment dynamics (Top simple decision-making task, bottom integration task). **(a)** Example input stimulus (green) and target output (red) time series for each task with noted periods. Amplitudes are shown in arbitrary units. The output is a constant binary value (± 1) determined by the stimulus, while the input varies with added noise (top) or is a random signal (bottom).**(b) W**^**in**^ alignment with the input peaks during signal period (0.25 − 0.50) for Simple DM and for signal and intermediate for Integration task, then decays post-decision in both. **(c) W**^**out**^ alignment with output activity (0.25 − 0.30) increases sharply during the decision period. Bars are colored as the corresponding time periods.

In contrast, output alignment (Fig. 6c) is minimal during the signal period (positive: 0.024 ± 0.004; negative: 0.024 ± 0.005) and increases during the intermediate (positive: 0.274 ± 0.008; negative: 0.276 ± 0.008) and decision periods (positive: 0.296 ± 0.000; negative: 0.299 ± 0.000). For the integration task, output alignment is also low during early periods (signal: positive 0.013 ± 0.004, negative 0.013 ± 0.004; intermediate: positive 0.021 ± 0.010, negative 0.027 ± 0.013) and increases sharply during the decision period (positive: 0.283 ± 0.041; negative: 0.298 ± 0.002).

These results quantitatively confirm a temporal segregation between sensory encoding and decision execution, with input-aligned representations dominating early epochs and output-aligned representations emerging at decision time.

This temporal decoupling was conserved across both reactive and integrative tasks. The low correlation between activity and **W**^out^ prior to decision onset is largely imposed by the task design and loss function, which enforce a near-zero target output during this period. Thus, low output alignment before the decision epoch primarily reflects the absence of a required readout signal rather than a failure of the network to engage decision-related dynamics. The relevant dynamical transition is the sharp increase in alignment at decision onset, marking the recruitment of the output mode.

### 3.4 Distributed Connectivity and Task Performance

Single-unit ablations markedly altered the network dynamics and impaired task performance (Fig. 7). We systematically evaluated the impacts of single-unit removals across trained networks and tasks, and found that eliminating even a single unit consistently impaired performance and altered the learned dynamics. Fig. 7 illustrates a representative example of this effect. The resulting trajectories deviated substantially from the intact network, indicating that the computation is not localized to specific units but instead emerges from the coordinated activity of the recurrent population. These findings suggest that the learned dynamical solution is highly distributed and sensitive to local perturbations.

**Fig. 7.**
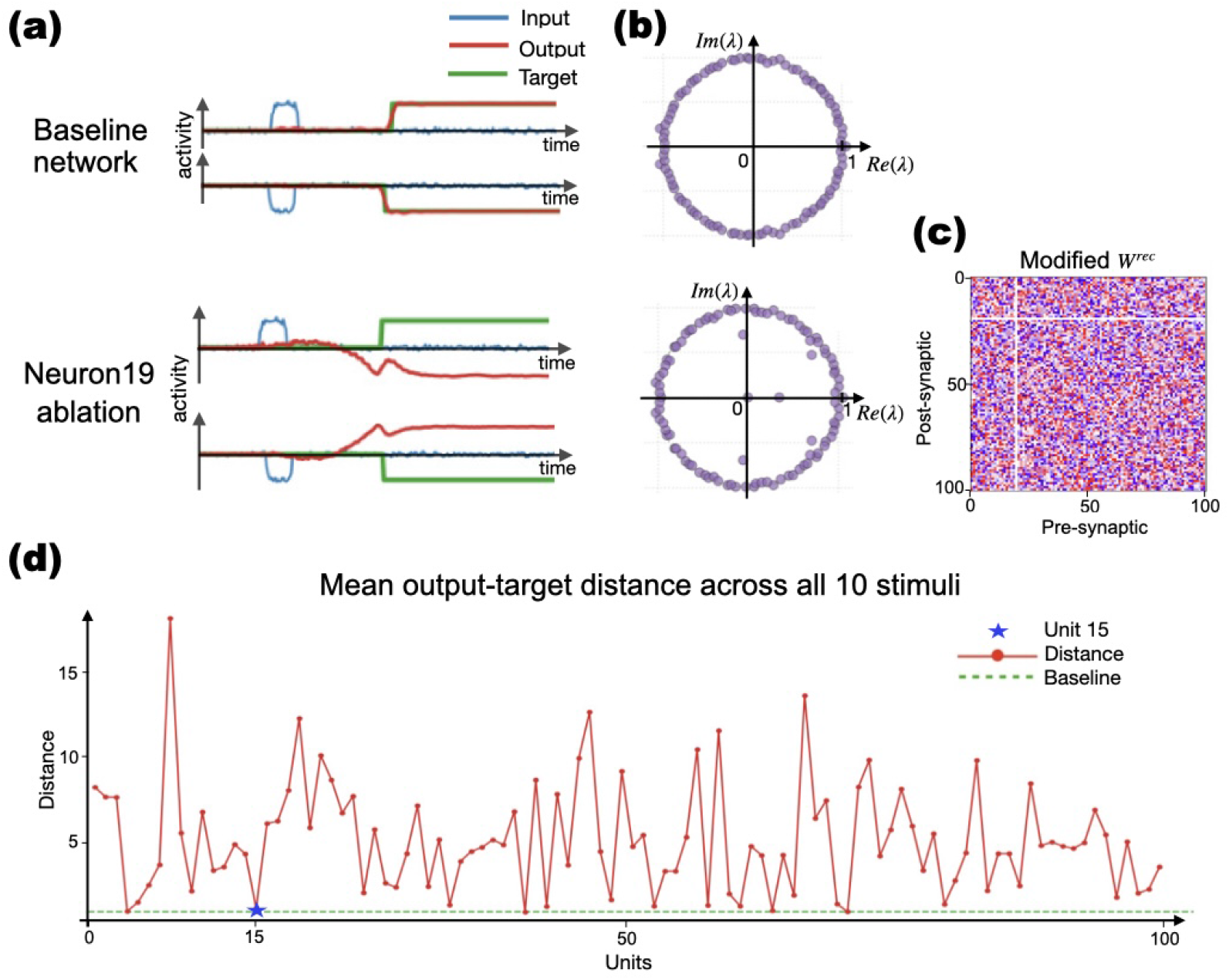
Unit ablation analysis after single-unit removal. **(a)** Baseline output (top) compared with output after removing one representative unit (bottom). **(b)** Eigenvalue spectrum of the intact (top) and ablated (bottom) network. **(c)** Recurrent weight matrix with the ablated unit’s row and column zeroed, illustrating the disconnection procedure. **(d)** Mean output-target distance across all 10 stimuli for each single-unit ablation (dashed line: 2.5 threshold). Most ablations exceed the threshold, indicating distributed coding; a small number of units (e.g., unit 15) can be removed with minimal performance loss.

Ablations also altered the recurrent eigenvalue distribution (Fig. 7b-c), with similar patterns across replicas and tasks as in Jarne and Caruso (2023). This alteration can be measured by output-target distance, which exceeded 2.5 for over 90% of ablated units (see Fig. 7d). The 2.5 threshold corresponds to the maximum target-output distance observed in well-trained intact networks when averaging over 10 pairwise sample distances of fixed length (comparing each stimulus target with its corresponding network output). Distances exceeding this value thus indicate deviations beyond the range of variability observed in properly trained networks. This measurement suggests that most single-unit ablations caused the network’s output to deviate substantially from its intended target across all tested stimuli (see SI animations). These disruptions not only affected responses to stimuli but also perturbed global network dynamics, indicating a distributed coding scheme.

Networks tolerated limited ablations of certain units (e.g., unit 15), whereas removal of most units (e.g., unit 19) substantially impaired performance (Fig. 7d). This heterogeneity indicates that recurrent units contribute unequally to the learned computation. Nevertheless, because performance deteriorated following ablation of the majority of units, the computation cannot be attributed to a small set of specialized neurons and instead appears to emerge from distributed population dynamics. We note that this ablation analysis characterizes the degree to which single units are individually dispensable, but does not by itself identify why particular units are more critical than others, for instance, whether this reflects their contribution to specific temporal dynamics.

### 3.5 Robust Temporal Processing Across Input Modalities

We next asked whether different input parameterizations, pulse amplitude vs. inter-pulse interval, lead to distinct network dynamics. Comparing the Multi-interval Amplitude- and Distance-based tasks (SI Fig. S1-S2), we found that dynamics were *quantitatively similar*: the distributions of the Sequentiality Index (SeqIdx) overlapped across tasks (Fig. 8a), and the low-dimensional PCA subspace was closely aligned (small principal angles between the top PC subspaces). Any between-task differences were of similar orders of magnitude when contrasted with the stimulus-dependent variability within a single network (see Fig. 8a), suggesting that input format does not strongly constrain the dynamical regime. We note that overlapping SeqIdx distributions indicate that both input parameterizations recruit comparable degrees of sequential versus persistent population activity, not that the underlying dynamics are identical.

**Fig. 8.**
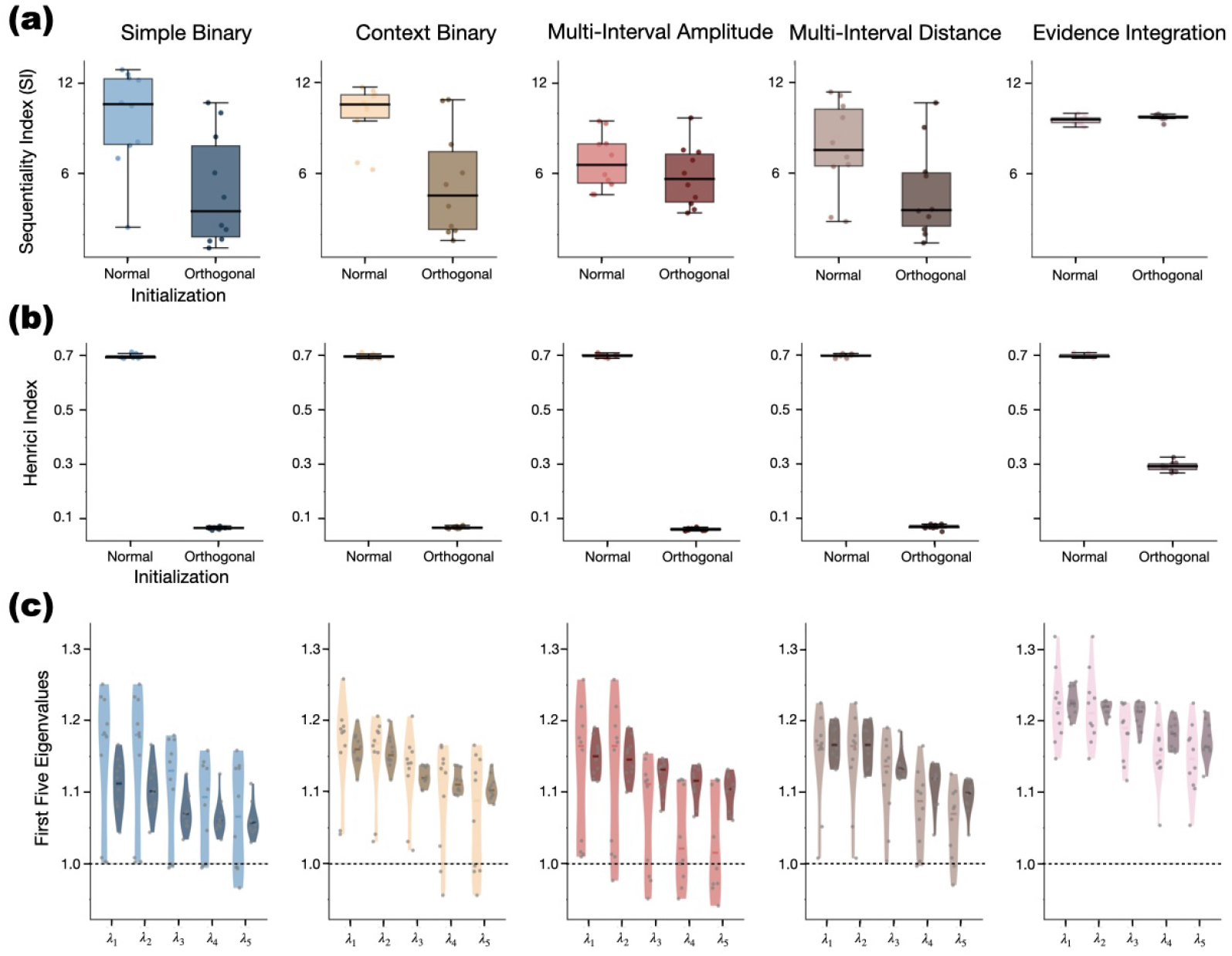
Sequentiality, non-normality, and eigenvalue magnitudes across tasks and initialization. (**a**) Sequentiality Index (SI) for each task, comparing Normal (lighter) and Orthogonal (darker) initialization. Boxplots show median and quartiles; dots represent individual replicas. (**b**) Henrici index (measure of non-normality). Orthogonal initialization consistently yields near-normal matrices (Henrici ≈ 0). (**c**) Top-5 eigenvalue moduli (|*λ*_1_| to |*λ*_5_ |) for each task, displayed as violins with median lines. Grey points show individual networks; the dashed line indicates the unit circle.

Although trajectory geometries differ across tasks in reduced-dimensional space, all trained networks shared non-normal recurrent structure. However, the dominant temporal dynamics varied across trained replicas, with some networks exhibiting oscillatory solutions and others showing predominantly non-oscillatory (ramping/decaying) dynamics. Task differences therefore appear to modulate the parametrization of these shared recurrent structures rather than inducing entirely distinct temporal processing mechanisms.

### 3.6 Non-Normal Dynamics and Spectral Signatures of Task Complexity

Across tasks, trained networks consistently showed non-zero Henrici’s parameters (*d*_*F*_ > 0; (Fig. 8b)), indicating non-normal recurrent connectivity. The magnitude of non-normality varied with initialization scheme. We note that Henrici’s index is a global scalar summary of the weight matrix and does not explicitly capture input- or state-dependent dynamics. Thus, the associations reported here between non-normality and task complexity are empirical and do not imply a direct mechanistic link.

In integration tasks, eigenvalues were distributed farther from the unit circle (Fig. 8c), supporting a broader range of temporal frequencies and slower decay rates. In the TICT task (SI Fig. S3), non-normal dynamics gave rise to activity modes that simultaneously supported time estimation and categorical choice. In both cases, non-normality is consistent with transient amplification of specific activity patterns such as timing pulses or evidence ramps that were selectively boosted before the network settled into a stable state. This mechanism is crucial for encoding and manipulating information during delays and the decision-making processes.

### 3.7 Low-Dimensional Trajectories Encode Decisions and Temporal Context

Projection onto the first three principal components revealed structured, low-dimensional trajectories that represented task variables. In our tasks, the first three principal components typically captured the majority of neural variance (e.g., >90% for the simple DM task; see SI Fig. S7-S10). PCA projections for other tasks (Figs. 3c, 4b) show qualitatively low-dimensional structure. To quantify precisely how different task conditions occupy distinct or shared population subspaces, we computed the principal angles between the leading three-dimensional PCA subspaces of trial-averaged hidden-state activity across task-relevant stages (pre-stimulus, stimulus, intermediate, and decision). This analysis was performed across multiple independently trained network replicas, with results reported as means ± SD.

In the simple delayed decision-making task, the principal angles between subspaces associated with positive and negative stimuli were large during the stimulus epoch (*PC*1 : 76.3° ± 11.8°, *PC*2 : 37.8° ± 13.8°, *PC*3 : 20.3° ± 12.4°), indicating that opposite-sign conditions drive population activity into substantially separated subspaces during stimulus presentation. During the intermediate (delay) period, angles remained moderately large for PC1 (53.0° ±18.4°), indicating partial separation. Notably, across the full trial, the subspaces nearly collapsed (*PC*1 : 2.4° ± 0.6°, *PC*2 : 1.2° ± 0.5°, *PC*3 : 0.0° ± 0.0°), consistent with the sign-flip symmetry of the population code: the full trial trajectory for positive and negative decisions occupies essentially the same low-dimensional subspace, related by a sign reversal. Explained variance was strongly dominated by the first PC for both conditions (positive: 91.3% ± 3.3%; negative: 93.0% ± 2.6%), confirming the highly low-dimensional character of the representation. The full quantification across epochs and replicas, including bar plots of the principal angles and the corresponding heatmap, is provided in SI Fig. S7 (panels b, c, d), SI Table S2 (explained variance), and the principal-angle Table S3.

In the windowed evidence integration task, the pattern of subspace separation was (*PC*1 : 41.0° ± 30.5°, *PC*2 : 30.5° ± 24.2°, *PC*3 : 4.9° ± 3.0°).During the intermediate period (*PC*1 : 81.7° ± 8.8°, *PC*2 : 40.5° ± 14.5°, *PC*3 : 10.4° ± 2.1°)) and was sustained at the decision epoch (*PC*1 : 43.2° ± 29.6°, *PC*2 : 11.7° ± 1.1°). The pre-stimulus angles were exactly 0° across all PCs, as expected given the identical initial state before any input is received. The first PC captured most of the variance in both conditions (*positive* : 91.0% ± 3.1%; *negative* : 77.0% ± 11.1%), again confirming highly low-dimensional dynamics. Complete visualization and numerical values are shown in SI Fig. S8 (panels b, c, d), SI Table S4 (explained variance), and the principal-angle SI Table S5. Together, these results indicate that the timing of subspace separation tracks whether task-relevant information is immediately available in the stimulus (as in the delayed binary task) or must be accumulated progressively (as in the integration task), providing a geometric signature of the underlying computational regime.

For the multi-interval tasks (decisions with multiple intervals, Category 2), the geometry reflects the additional demand of encoding one of several possible delay durations. In the amplitude-coded multi-interval task, distinct amplitude conditions occupied partially separated subspaces even during the intermediate period, with PC1 angles between the reference condition (*A*1+) and the longest delay (*A*8+) reaching 85.6° ± 7.1° during the intermediate phase (SI Fig. S9, panels a,b; explained-variance Table S6 and the three principal-angle in SI Tables S7, S8 and S9), indicating near-orthogonal representations for the most temporally distant conditions. The intermediate period showed consistently large angles across all amplitude pairs (PC1 angles ranging from 77° to 89°), while the decision period showed partial convergence. Across intervals, the explained variance in PC1 decreased progressively with amplitude (from 95.2% ± 0.7% for *A*1+ to 71.3% ± 5.9% for *A*8+), reflecting the greater complexity of dynamics required to encode longer delays. A similar pattern of subspace separation during the intermediate period was observed in the distance-based multi-interval task(SI Fig. S10, explained-variance in SI Table S10 and three principal-angle in SI Tables S11, S12 and S13), with near-orthogonal intermediate-period subspaces for temporally distant conditions (PC1 angles up to 77° between *T*_20_ and *T*_80_ at the intermediate stage). These results show that the network encodes different delay durations in partially orthogonal subspaces during the delay period itself, and that this orthogonality is largely resolved at decision time, consistent with a shared decision-axis readout irrespective of the preceding interval.

### 3.8 Spectral Features and the Dynamical Regime

Spectral structure varied systematically with task demands, consistent with prior work linking spectral geometry to cognitive dynamics (Jarne 2021, 2022; Jarne and Laje 2023). Outlier eigenvalues which emerged during training were associated with network dynamics. Tasks involving integration displayed eigenvalues farther from the unit circle and broader spectral dispersion (Fig. 8c), consistent with the need for richer temporal structure. The spectral patterns were qualitatively similar across initialization schemes: both random normal and orthogonal initializations produced comparable eigenvalue dispersion and outlier structure after training.

We asked whether the spectral structure of the trained recurrent weights varied systematically with task demands. To address this, we compared the eigenvalue spectra of *W* ^Rec^ across tasks of increasing complexity (simple DM, multi-interval, and integration). Tasks requiring richer temporal structure (e.g., integration over variable windows) produced eigenvalue distributions with greater dispersion and outlier eigenvalues farther from the unit circle (Fig. 8c), consistent with the need for a broader range of characteristic timescales. We emphasize that these are empirical associations; as noted in Section 2.5, eigenvalues of *W* ^Rec^ alone do not determine the effective dynamics of a nonlinear, input-driven network.

This pattern was confirmed statistically: a Kruskal-Wallis test on the top-5 eigenvalue moduli across tasks was significant at every rank (*λ*_1_–*λ*_5_; all *p* < 0.0001). Dunn’s post-hoc tests with Bonferroni correction showed that the Windowed Evidence Integration task differed significantly from every other task at every rank (all *p* < 0.005), whereas no pairwise differences were significant among Simple DM, Context-dependent DM, Multi-interval Amplitude-based, and Multi-interval Distance-based tasks. This indicates that the increased spectral dispersion is specifically associated with the integration task rather than reflecting a smooth gradient with task complexity across all tasks studied.

These results together show that similar task demands can be implemented through different dynamical regimes, including both oscillatory and non-oscillatory solutions, with varying degrees of alignment to the readout dimension during task. While specific features depend on task parametrization, the underlying computational strategies exhibit a common structure. Across tasks, these analyses reveal that temporal representations arise from a shared set of dynamical features, modulated by task parametrization rather than determined by it.

We next asked whether oscillatory and non-oscillatory networks could be discriminated using spectral descriptors extracted from the final recurrent connectivity. A logistic regression trained on these descriptors, with regularization strength selected by nested cross-validation and evaluated by leave-one-out prediction, discriminated the two classes above a task-stratified permutation null (balanced accuracy 0.68, *ROC* − *AUC*0.67, *p* = 0.005; Fig. 9b). This is notable given that no individual descriptor was significantly associated with the regime label after FDR correction (Fig. 9a). Thus the discrimination arises from the combination of weakly informative features rather than from any single one. Discrimination was modest, and these descriptors are a coarse characterization of a 100 × 100 recurrent matrix, so this result indicates that the final connectivity retains some regime-related spectral information, without implying that the regime is strongly determined by it. The complete correlation structure among all spectral descriptors, behavioural metrics, initialization scheme, and task complexity is presented in the hierarchically clustered correlation matrix (SI Fig. S5).

**Fig. 9.**
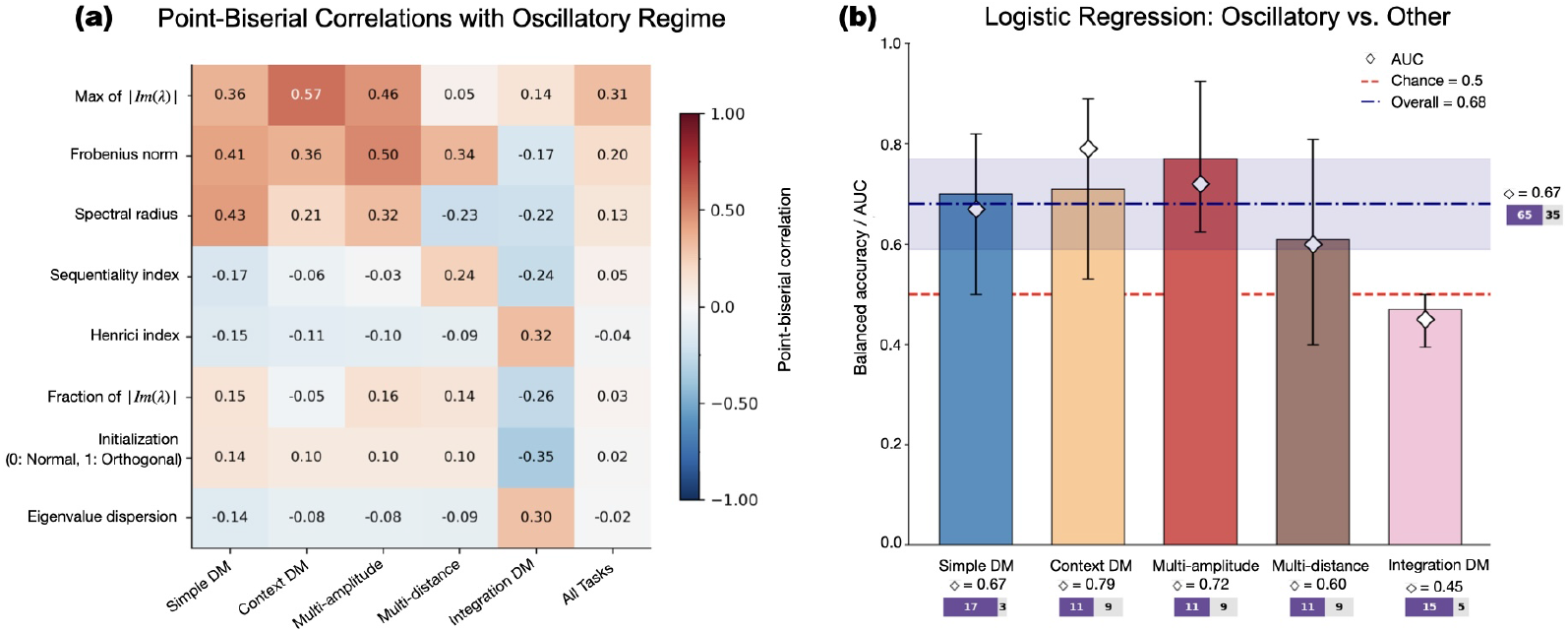
Regime predictability from final connectivity. (**a**) Point-biserial correlation between spectral features (and SI) and the oscillatory/non-oscillatory label, calculated per task and overall (ALL TASKS). Rows are sorted by the absolute correlation strength in the ALL TASKS column. After applying permutation test + FDR correction (*N*_*P ERM*_ = 500, Benjamini-Hochberg correction), none of the individual magnitudes correlates significantly either for the networks separated by training of the individual tasks or for all the unseparated tasks. After correcting the p-values, none of the correlations remains significant (represented by the absence of asterisks in the heat map) (**b**) Leave-one-out cross-validated accuracy of a logistic regression classifier that discriminates oscillatory from other trained networks using spectral descriptors extracted from the final recurrent connectivity, together with the SeqIdx and initialization scheme. Coloured bars show task-specific accuracies, with diamonds indicating the corresponding AUC. The horizontal dashed line indicates chance performance (0.5), and the navy line denotes the overall accuracy with a 95% bootstrap confidence interval (percentile method, 2000 resamples). A task-stratified permutation test comparing the observed overall balanced accuracy to a null distribution (not a binomial test) indicated that the overall classifier performs significantly above chance (p=0.005). Note that this test applies only to the overall classifier and not to the individual task bars. The purple/gray mini-bar shows the number of used oscillatory and others.

Then, we directly tested whether oscillatory and non-oscillatory trained networks could be discriminated using spectral descriptors extracted from the final recurrent connectivity. A logistic regression trained on spectral features (max imaginary part, Henrici index, initialization, etc.) and validated with a leave-one-out procedure classifies oscillatory vs. non-oscillatory networks significantly above chance (Fig. 9b). This holds overall and for four of the five task families; the Windowed Evidence Integration task is the exception, which follows from the class imbalance in Table 2, where 15 of 20 replicas are oscillatory and none are purely ramping, leaving too few non-oscillatory examples for a stable balanced-accuracy estimate. These results indicate that the final recurrent connectivity contains sufficient spectral information to discriminate oscillatory from other trained networks using a small set of spectral descriptors. When networks are grouped according to their final dynamical regime, oscillatory networks consistently exhibit higher maximum imaginary part and Oscillation Index, while mixed networks span a wide range of Sequentiality Index (SeqIdx), values (SI Fig. S6).

### 3.9 Generalization Analysis

To assess generalization, we evaluated trained networks in the multi-interval amplitude-based task and the windowed evidence-integration task. We considered both intermediate parameter values within the training range (interpolation) and values outside the trained regime (extrapolation).

Network performance was assessed by generating novel test trials at each parameter value (20–40 trials per value) and evaluating the trained RNNs on three metrics. Accuracy was defined as the fraction of trials for which the sign of the mean network output during the last 25 time steps matched the target sign; mean squared error (MSE) was computed over the same interval and only for trials with a non-zero target. To quantify temporal precision, we detected the response onset as the first time step where the output exceeded a threshold of 0.3 with the correct sign, and computed the relative onset error as the percentage deviation of the predicted onset from the target onset, averaged across all valid trials. All metrics were aggregated across network replicas and reported as mean ± standard deviation.

Across replicas, in the multi-interval amplitude-based task, interpolation to intermediate values within the training range was accurate, and extrapolation beyond that range was asymmetric. Accuracy remained comparatively high for amplitudes above the trained range (0.920.14 and 0.840.29 for Normal and Orthogonal initialization, respectively), though accompanied by a systematic negative timing bias, whereas accuracy dropped sharply below the trained range (0.740.28 and 0.530.23). In contrast, in the windowed evidence-integration task, networks trained with a single 200-step integration window exhibited robust extrapolation across a wide range of window lengths (50-430 steps). In contrast, in the windowed evidence-integration task, networks trained with a single 200-step integration window exhibited robust extrapolation across a wide range of window lengths (50-430 steps). For more details, see Supplementary Information Section 4.3, SI Fig. S11 and SI Fig. S12.

## 4 Discussion

Our findings demonstrate that RNNs trained on temporally parameterized decision-making tasks develop low-dimensional dynamics and a variety of functionally equivalent solutions and show that sign-flip equivariance imposed by zero-bias initialization and odd nonlinearities is preserved exactly during training on symmetric tasks, despite noisy inputs and stochastic optimization. These findings reflect broader principles of neural computation observed in biological and artificial systems, while also revealing how task complexity and network connectivity shape distinct learning regimes.

In particular, we find that eigenvalue dispersion reflects task demands, suggesting a link between dynamical structure and computational load. Specifically, the Windowed Evidence Integration task consistently produced broader spectral dispersion and slower dynamical modes than all other tasks, whereas multi-interval tasks did not differ significantly from simple delayed decision-making tasks in this respect, suggesting that the demand for continuous temporal integration, rather than task complexity more broadly, is what recruits a wider range of effective timescales.

The preserved sign-flip equivariance produced mirrored population trajectories and anti-correlated unit responses reminiscent of population-level “anti-neuron” motifs, similar to Yang et al. (2019), where RNNs trained on multi-task paradigms develop shared, reusable dynamical motifs. Similarly, our observation that decision trajectories cluster in low-dimensional subspaces mirrors the “untangled” population responses observed in motor cortex (Russo et al. 2018). However, unlike these studies, here the preserved sign-flip equivariance emerged from the combination of zero-bias initialization, odd nonlinearities, and sign-symmetric training.

Across tasks, the relative prevalence of dynamical regimes varied (Table 2), indicating that task parameterization influences which of several computational strategies is more frequently adopted. We do not attribute this variation to a specific task dimension. Our tasks differ simultaneously in delay structure, input format, integration demand, and number of intervals, and the differences in prevalence were not stable across reasonable choices of the analysis window used to compute the activity metrics. We therefore report these differences descriptively, and leave the identification of the responsible task variable to a study designed to manipulate it directly.

Several recent studies have used task-trained RNNs to characterize solution diversity and dynamical motifs. Orhan and Ma (2019) asked what conditions favour sequential versus persistent short-term memory representations by sweeping circuit hyperparameters. Driscoll et al. (2024) identified compositionally reusable dynamical motifs in multitask-trained networks, emphasizing cross-task generalization via overlapping neural subspaces. Turner et al. (2021) developed algorithmic tools for charting the RNN solution space. Our work addresses a complementary question: how do networks jointly encode elapsed time and decision variables when the temporal demands of the task are parametrically varied within a fixed training framework? Rather than multitasking flexibility, we focus on degeneracy in single-task solutions and find: (i) predominantly oscillatory, ramping/decaying, and mixed dynamical regimes achieving comparable performance, while remaining distinguishable through task-specific spectral signatures and antisymmetric stimulus-response mappings, (ii) a temporally structured recruitment of the readout-aligned mode at decision time, consistent with the task design and output constraints imposed during training, and (iii) preservation of near-sign-flip equivariance in odd-symmetric (tanh()) architectures through training.

The degeneracy of solutions, meaning that distinct networks achieve similar performance through oscillatory or non-oscillatory dynamics, aligns with Huang et al. (2024), who showed that RNNs exploit diverse dynamical motifs to solve identical tasks. Our networks exhibited a spectrum of dynamics with Sequentiality Indices (SI) values ranging from persistent activity to periodic oscillations, where persistent activity (low SI) and periodic dynamics (high SI) coexist, (see Fig 8). However, Huang et al. (2024) define complexity through loss landscape roughness, whereas our tasks inherently modulate complexity via temporal integration requirements (e.g., multi-interval vs. cued decisions). We suggest that task structure, not just optimization dynamics, critically influences solution diversity.

The coexistence of oscillatory and ramping/decaying dynamics within a single task architecture is also relevant for the time-perception literature. Neurophysiological studies have reported both ramping activity in prefrontal and parietal cortex (Merchant et al. 2013b; Mello et al. 2015) and oscillatory or sequential patterns in the striatum and hippocampus (Matell and Meck 2004) during interval-timing tasks. Theoretical work has shown that recurrent networks can intrinsically encode time through state-dependent trajectories that are read out by a downstream observer (Buonomano and Laje 2010), and that the shape of these trajectories (ramp vs. oscillation) may reflect distinct but interchangeable timing mechanisms. The solution degeneracy we observe mirrors this biological diversity and suggests that both regimes can serve as functionally equivalent temporal integrators. In particular, ramping dynamics naturally support scalar variability when coupled with a threshold (Simen et al. 2016), while oscillatory “cycle-counting” can implement precise, discretised timing (Matell and Meck 2004).

In the multi-interval amplitude-based task, trained networks interpolated within the training range, but extrapolation beyond it was asymmetric: performance degraded only modestly for amplitudes above the trained range, while it collapsed for amplitudes below it. Networks trained on the windowed evidence-integration task, by contrast, exhibited robust extrapolation across a wide range of integration windows in both directions (Valente et al. 2022). This pattern suggests that the learned amplitude-to-delay mapping is not simply confined to the trained interval, but is more robust in the direction of longer, previously unseen delays than in the direction of shorter ones, whereas the integration task’s solution is close to scale-invariant in both directions because the underlying computation reduces to simple accumulation.

Our networks exhibit signatures of substantial recurrent reconfiguration and therefore do not appear consistent with a purely lazy-learning regime. In the lazy learning regime, networks remain close to their initialization and rely primarily on the readout layer to solve the task (Liu et al. 2023). In contrast, training in our models produced substantial changes in the recurrent weights, accompanied by broad spectral reorganization and the emergence of distributed recurrent activity patterns, indicating that the learned solutions deviated considerably from their initial state. Consistent with this reconfiguration, the aggregate marginal distributions of **W**^in^, **W**^Rec^, and **W**^out^ after training did not differ substantially in overall shape from their initialization distributions; however, the specific configuration of individual weights, and consequently the eigenvalue spectrum of **W**^Rec^, changed substantially during training, indicating that training reorganizes connectivity at the level of individual weights and their collective spectral structure rather than at the level of the population-level weight distribution.

At the same time, the alignment between recurrent activity and the readout vector remained weak during the delay/integration epoch and increased sharply near decision onset. Because the target output is constrained to remain near zero before the response period, weak early alignment with **W**^out^ is expected from the task design itself. The relevant dynamical feature is therefore the temporally localized recruitment of readout-aligned activity during the decision epoch, consistent with low-dimensional readout-aligned representations observed in trained recurrent networks (Braun et al. 2022).

After training, readout alignment was weak during integration (*r* ≈ 0) and increased near decision time (*r* ≈ 0.25–0.30), suggesting that task-relevant information is initially encoded in distributed recurrent dynamics before becoming progressively aligned with the behavioural readout dimension. These results indicate that tasks requiring delayed or temporally integrated decisions can involve substantial recurrent reconfiguration during learning while still producing temporally selective readout-aligned activity at inference.

The distributed nature of the learned solutions, as evidenced by the substantial performance decrements with single-unit ablations, echoes principles of degeneracy in biological systems (Edelman and Gally 2001). Unlike the specialized circuits in Mante et al. (2013), our networks relied on preserved sign-flip equivariance. This distributed coding aligns with Stringer et al. (2019)’s observation that high-dimensional neural populations encode stimuli uniformly, but contrasts with modular architectures in Yang and Wang (2020). Notably, our finding that PCA trajectories separate cue-dependent and cue-independent states (SI Fig. S4) parallels how prefrontal cortex gates task-relevant information (Russo et al. 2020), suggesting RNNs recapitulate biological gating mechanisms through dynamical (not structural) modularity.

While Huang et al. (2024) defines complexity through the loss landscape’s geometry, our work operationalizes complexity via task demands. For instance, multi-interval tasks required networks to bind temporal information to stimulus identity, increasing computational load compared to binary decisions. This complexity also manifested in eigenvalue spectra, where the task demanding continuous temporal integration produced broader eigenvalue distributions than the delayed decision-making tasks, akin to Jarne (2022).

Beyond eigenvalue dispersion, trained networks consistently exhibited non-normal recurrent connectivity (*d*_*F*_ > 0; Fig. 8b), regardless of task or initialization scheme. Non-normality is a necessary, though not sufficient, condition for transient amplification in linear systems (Bondanelli and Ostojic 2020), and has been associated with selective boosting of specific activity patterns, such as timing pulses or evidence ramps, before the network settles into a stable state (Asllani et al. 2018). We emphasize, however, that Henrici’s index is a global, connectivity-level descriptor and does not by itself establish that transient amplification occurs along the specific, input-driven trajectories realized during task execution; a full mechanistic account would require analysis of the state-dependent Jacobian along trial trajectories (Schuessler et al. 2020; Krause et al. 2022), which we leave for future work.

An open question is whether the dynamical regime ultimately adopted by a network is already constrained before training converges or instead emerges during optimization. Our results neither confirm nor exclude either possibility. Rather, they show that the final recurrent connectivity retains enough information to distinguish oscillatory from non-oscillatory solutions above chance, suggesting that the optimization process leaves identifiable signatures in the learned connectivity. Determining whether those signatures are already present before convergence would require analysis of the initial connectivity or intermediate training checkpoints.

For temporal tasks requiring internal clocks, recurrent weight adjustments (rich dynamics) appear unavoidable, even with high-rank initializations. This complements Braun et al. (2022)’s finding that temporal integration necessitates non-lazy solutions.

Together, the observed non-normal connectivity, delayed readout alignment, and evolving population geometry suggest that temporal computations are implemented through distributed transient dynamics rather than persistent activity confined to a fixed readout axis.

Discussion of the relevance between trained RNNs and biological systems requires addressing some limitations of the present study. First, our networks are small (N = 100 units) and fully connected, whereas some biological circuits are sparse and modular. Whether the dynamical regimes identified here scale to larger, sparser architectures remains open. Second, our generalization analysis reveals that extrapolation is direction-dependent in the multi-interval amplitude-based task, holding up reasonably well for longer, untrained delays but collapsing for shorter ones, suggesting that learned solutions are only partially tied to the specific trajectories seen during training rather than encoding a fully abstract timing rule. Future work should examine whether architectural constraints (e.g., Dale’s law, sparse connectivity) may promote more generalizable temporal representations. Third, the present tasks employ discrete response epochs; extending this framework to continuous or naturalistic timing demands will be critical for bridging the gap between trained RNNs and biological systems. Fourth, our regime-predictability analysis (Fig. 9) was based on ensembles of ten trained replicas per task and initialization condition; while sufficient to detect above-chance discriminability of the oscillatory regime from the final recurrent connectivity, a substantially larger ensemble would be needed to disentangle the causal contributions of initialization, task parameterization, and stochastic optimization to the emergence of a given dynamical regime. A complementary approach to establish causality would be to pre-train networks on a task that strongly favours one dynamical regime (e.g., a fixed-delay task biased toward oscillatory solutions) and then continue training on a task with more neutral temporal demands, to test whether the initially acquired regime persists or is overwritten; we leave both directions for future work. Finally, whether similar degeneracy and symmetry properties persist in architectures with gating, sparsity, or biological constraints remains an open question.

Taken together, our findings suggest that elapsed time can be represented through multiple recurrent dynamical implementations that nevertheless converge to similar low-dimensional computational representations.

## Conclusion

Our results underscore the interplay between task structure, initial connectivity, and learning regimes in shaping RNN dynamics. Building on concepts from solution degeneracy, population geometry, and biological computation, this work advances our understanding of how artificial and biological networks balance flexibility and efficiency in temporal decision-making. Future work will investigate how these principles –degenerate dynamics, population-wide symmetry, and temporal decoupling – scale to more ethologically relevant tasks, and whether similar signatures exist in biological neural recordings during flexible time-dependent decisions.

## Supplementary information

We developed our code in Python based on the TensorFlow (Abadi et al. 2015) and Keras (Chollet et al. 2015) frameworks to train the networks for each task and to perform analysis. The code and trained networks (saved in HDF5 format) can be accessed at https://github.com/katejarne/RNNs for DM and time representation

## Acknowledgements

C.J. was supported by PICT-2020-01413 and a UNQ project (EXPTE.1520/25). Z.P.K. and K.J. were supported by the NSF (CRCNS program; DMS-2207700). KJ was additionally supported by the NIH (R01 MH130416). K.J. and R.Y. were supported by the NSF (NeuroNex program DBI-1707400). T.L.E. was supported by the NIH (1K99NS127855-01A1).

## Ethical Approval

Not applicable.

## Authors’ contributions

C.J. developed the code, performed simulations, analyzed data, and wrote the manuscript. Z.K. and K.J. supervised research and suggested analysis of neural trajectories and network studies. R.Y. edited figure visualizations. Z.K., K.J., T.E., R.Y., and C.J. equally contributed to manuscript discussion, writing, and editing.

