## Supplementary material for "Task-Parametrized Dynamics: Representation of Time and Decisions in Recurrent Neural Networks": SI pdf

### 1 Supplementary figures for tasks from Table 1

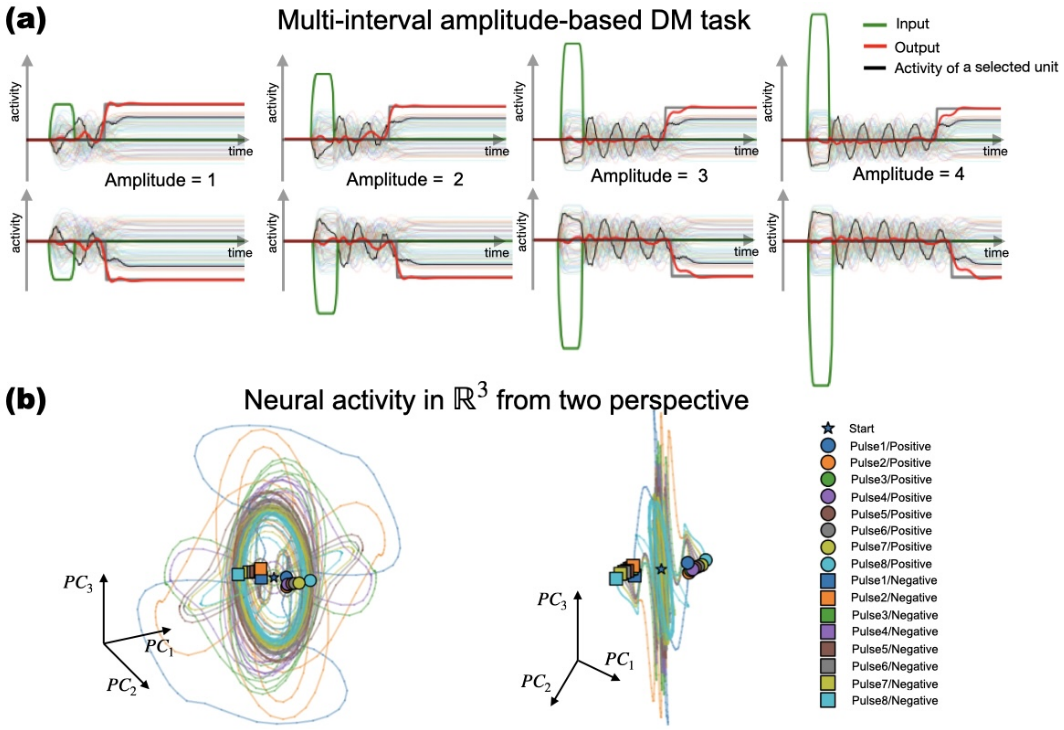


Figure S1: Multi-interval Amplitude-based Decision Making task. It uses eight distinct stimulus amplitudes, each corresponding to a different delay time for target responses. While the output remains binary (positive or negative), the network must store the appropriate response interval based on the stimulus's multiple amplitudes. Panel a) displays four time series with a positive stimulus of varying amplitudes, as well as the network's response to the same stimulus when it is negative. The input signal is represented in green, the output value in red, and the activity of one randomly selected neuron (which is consistent across all series) is highlighted in black. Panel b) illustrates the activity in reduced-dimensionality space (PCA) from two different perspectives, superimposing all possible response time intervals for which the network was trained.

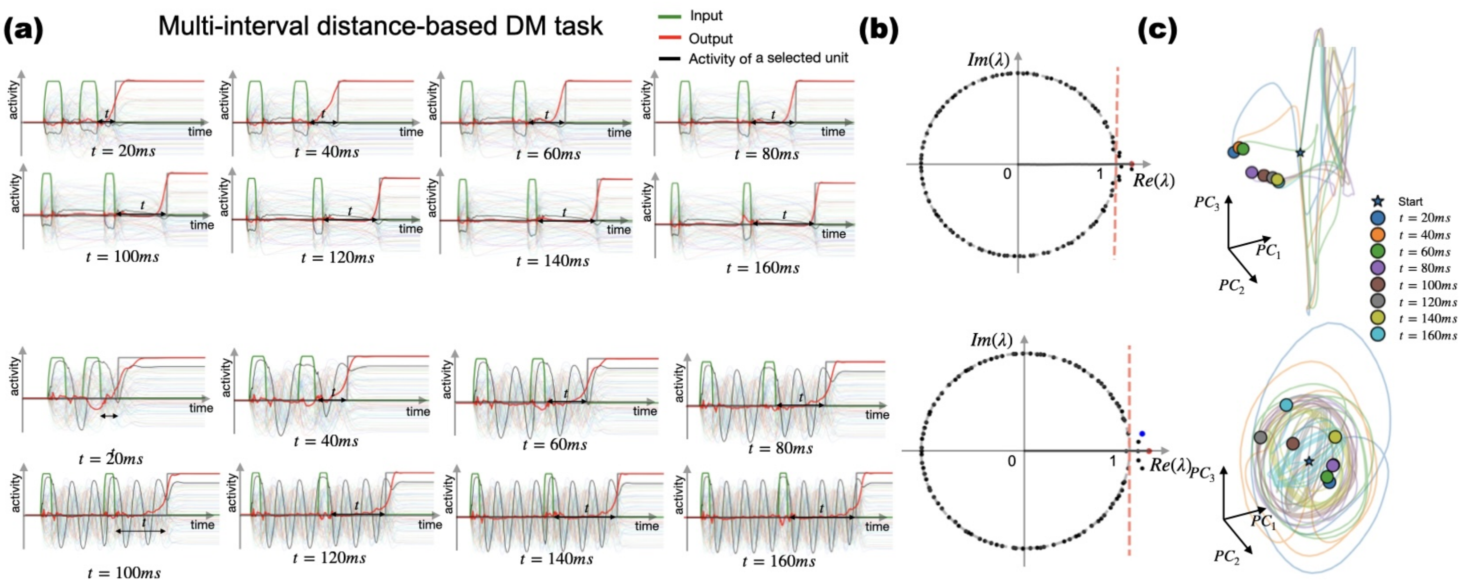

Figure S2: Multi-interval Distance-based Decision Making task. The response time is encoded as the time between two consecutive input pulses. We compared two different networks trained to perform the same task: the Top network and the Bottom network. Column a) presents the eigenvalue decomposition of each trained network. Column b) illustrates the internal behaviour across the eight intervals, highlighting that the Bottom network is primarily characterized by oscillation. Column c) displays the differences in trajectories within the PCA space, showing that different temporal intervals converge at neighbouring points in the PCA representation for each trained network. 1 ms is equal to 1 time step.

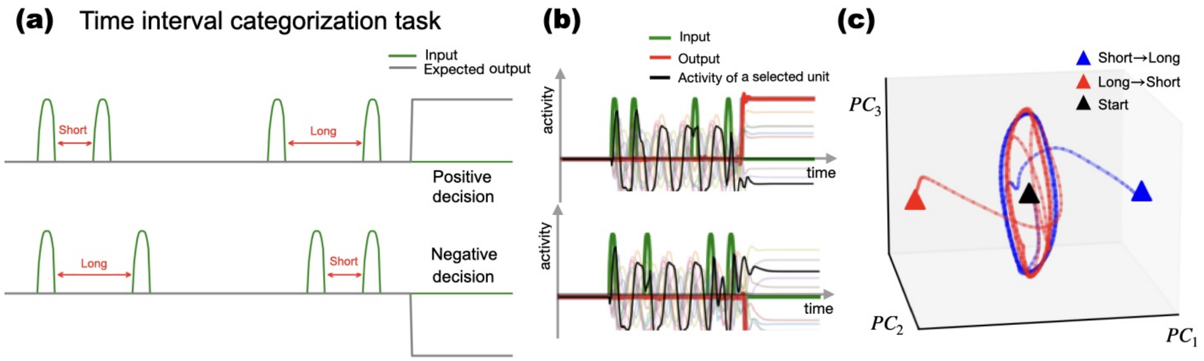

Figure S3: Time-Interval Comparison Task (TICT) dynamics. (a) Task Description. (b) Unit activity and (c) PCA trajectory (bottom), consistent with oscillatory dynamics encoding time intervals through repeated cycles.

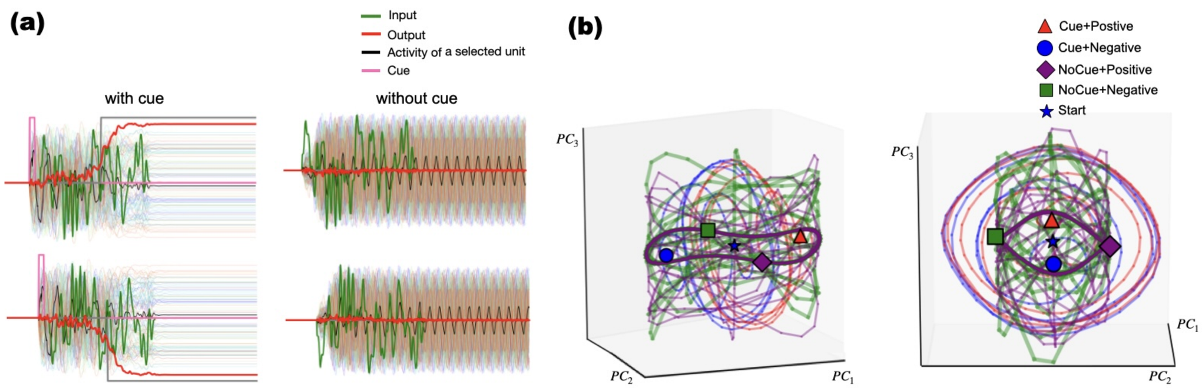


Figure S4: Cued Integration time Decision Making (time series panel a), and PCA panel b): orthogonal PCA planes for cue-present (decision axis) vs. cue-absent (null axis).

#### 2 Supplementary Animations

Files with SI animations are available in:
https://github.com/katejarne/RNNs_for_DM_and_time_representation/tree/main/Animations_compressed

#### 3 Supplementary Methods

All results reported in the main text correspond to bias-free networks (bias = 0). Configurations with trainable biases are included here only if the reader wants to perform a comparison, and for main text Section 3.2.

Table S1: Hyperparameters and training configuration used for the recurrent neural network across all decision-making tasks. The table reports settings that ensure reproducibility, including weight initialization distributions, optimizer parameters, loss function, dataset sizes, and early stopping criteria.

| **Parameter** | **Value** | **Notes** |
|---|---|---|
| | *Weight initialization* | |
| Input kernel | `Gaussian` | Default for SimpleRNN |
| Recurrent bias | Zeros | Enabled (use_bias=True or False) |
| Recurrent kernel | Orthogonal/RandomNormal | Base: RandomNormal(mean=0, stddev=$\sqrt{1/N_{rec}}$) |
| Dense kernel | `glorot_uniform` | Default for Dense layer |
| Dense bias | Zeros | Enabled/or not |
| | *Optimizer* | |
| Type | Adam | |
| Learning rate | 0.0001 | |
| $\beta_1$, $\beta_2$ | 0.9, 0.999 | |
| $\epsilon$ | $10^{-8}$ | |
| Gradient clipping | $\ell_2$ norm = 1.0 | clipnorm=1.0 |
| Loss function | Mean Squared Error (MSE) | |
| | *Dataset and training* | |
| Batch size | 64 | |
| Epochs (max) | 20 -40 | |
| Training samples | $N_{\text{train}}$ = sample_size − 50 | First 50 trials omitted from training |
| Total dataset size per task | `sample_size` depends on task:<br>Simple DM family: 15050<br>Integration task family: 90300 (6×15050)<br>Multi Amplitude: 90300<br>Interval comparison: 15050 | |
| | *Early stopping* | |
| Metric | Training loss | |
| Threshold | $5 \times 10^{-5}$ | Stops when loss < threshold |
| Patience | None | Immediate stop at first epoch below threshold |
| | *Other* | |
| Recurrent constraint | None | |
| Bias usage | True/False | For both RNN and Dense layers |
| Model checkpointing | Saved every epoch | Format: .keras, monitor='val_loss' |

#### 3.1 Sequentiality Index computation

Sequentiality was quantified following the framework introduced by [1]. For each trial, recurrent population activity was extracted from the hidden layer of the trained RNN, yielding an activity matrix:

$$H = \{h_i(t)\} \in \mathbb{R}^{N \times T} \tag{1}$$

where $N$ denotes the number of recurrent units and $T$ the number of time steps. To exclude silent neurons, only units with mean activity above a fixed threshold were included:

$$\frac{1}{T}\sum_{t=1}^{T} h_i(t) > r_{\text{threshold}}, \tag{2}$$

with $r_{\text{threshold}} = 0.1$.

For each selected unit $i$, the peak time was defined as

$$t_i^* = \arg\max_t h_i(t). \tag{3}$$

For each unit, activity within a temporal window of fixed size ($w = 5$ time steps) that was centered around the peak was compared to activity outside this window. The ridge-to-background ratio was defined as

$$R_i = \log\left(\frac{\frac{1}{|W_i|}\sum_{t\in W_i} h_i(t)}{\frac{1}{T-|W_i|}\sum_{t\notin W_i} h_i(t)}\right), \tag{4}$$

where $W_i$ denotes the set of time indices within the peak-centered window. The ridge component was computed as the average across active units:

$$R = \frac{1}{N_{\text{active}}}\sum_i R_i. \tag{5}$$

The entropy of the distribution of peak times $\{t_i^*\}$ was computed using a histogram with 20 bins. Let $p_k$ denote the normalized histogram counts (after adding a small smoothing constant $\epsilon = 0.1$ to avoid numerical instability). Entropy was defined as

$$S = -\sum_k p_k \log p_k. \tag{6}$$

Entropy was not normalized by log(bins); therefore, absolute SeqIdx values are not directly comparable in scale to those reported in the original study, although the computational principle is preserved.

The Sequentiality Index was defined as the sum of the ridge and entropy components:

$$\text{SeqIdx} = R + S. \tag{7}$$

SeqIdx was computed independently for each trial.

##### 3.2 Regime Classification Procedure

**Spectral criterion (Oscillation Index).** For each trained network, we extracted the recurrent weight matrix $\mathbf{W}^{\text{Rec}}$ and computed its eigenvalues. The Oscillation Index was defined as $\text{OI} = |\,\text{Im}(\lambda^*)|/|\lambda^*|$, where $\lambda^*$ is the eigenvalue with the largest modulus. A network was classified spectrally as *Oscillatory* if $\text{OI} \geq 0.15$, as *Mixed* if $0.05 \leq \text{OI} < 0.15$, and as *Ramping/Decaying* otherwise.

**Activity criterion and dual classification.** Activity-based classification used a fixed post-pulse window starting immediately after the input pulse offset and during the delay epoch. We used a smoothed square pulse (Hann-windowed), with Gaussian noise (5%, versus 10% during training) for this probe because the classification is designed to isolate the network's autonomous dynamics during the delay epoch, rather than to evaluate task performance under training conditions. A rectangular pulse introduces broadband transient energy at stimulus onset/offset that can contaminate the delay-window spectral estimate, and higher input noise increases the variance of the extracted PC1 trajectory, both of which can bias the peak-to-background ratio independently of the network's intrinsic regime. Reducing both sources of measurement noise yields a cleaner estimate of the dominant delay-period dynamics. To avoid cancellation by sign-flip symmetry, only trials with positive-polarity input were averaged. From the averaged trajectory, the first principal component (PC1) of the delay-window activity was extracted, linearly detrended, and its power spectral density was estimated via Welch's method. The peak-to-background ratio was computed as $\text{PBR} = \max_{f>0.10\,\text{cyc/step}} P(f) \,/\, \text{median}_{f>0.10\,\text{cyc/step}} P(f)$, and the $\log_{10}$ value was taken. A ramp index was defined as the absolute Pearson correlation between PC1 and time over the same window. Activity classification assigned *Oscillatory* when $\log_{10}(\text{PBR}) \geq 1$, else *Ramping/Decaying* if ramp index $\geq 0.1$, else *Mixed*. The final dual-consensus regime required both spectral and activity criteria to agree for an *Oscillatory* or *Ramping/Decaying* label; all other cases were labelled *Mixed*. The same thresholds were used for all tasks and initializations.

##### 3.3 Statistical analyses of regime predictability

To assess whether the dynamical regime could be inferred from the trained recurrent weights, we performed point-biserial correlations between spectral features and the oscillatory/non-oscillatory label, followed by a permutation test ($N = 500$ permutations) with Benjamini–Hochberg false discovery rate (FDR) correction across features and tasks. We further trained a logistic regression classifier on spectral descriptors (spectral radius, maximum imaginary part, eigenvalue dispersion, Henrici index, Frobenius norm, imaginary fraction, initialization scheme, and optionally the Sequentiality Index) to discriminate oscillatory from other networks. Regularization strength was selected via nested stratified cross-validation, and out-of-sample performance was evaluated with leave-one-out cross-validation. Overall balanced accuracy was compared against a task-stratified permutation null distribution. Differences in eigenvalue magnitudes across tasks were tested with Kruskal–Wallis and post-hoc Dunn tests. All analyses were implemented using scikit-learn and custom Python code.

##### 3.4 Generalization assessment

To assess how well the learned computations extended beyond the training conditions, we evaluated trained networks on novel parameter values both within the training range (interpolation) and outside it (extrapolation). For each parameter, 20–40 test trials were generated. Accuracy was defined as the fraction of trials in which the sign of the mean network output over the final 25 time steps matched the target sign. Mean squared error was computed over the same interval for trials with a non-zero target. Response onset was detected as the first time step where the output crossed a threshold with the correct sign, and the relative onset error $(t_{\text{pred}} - t_{\text{target}})/t_{\text{target}} \times 100$ was averaged across valid trials. All metrics were aggregated across network replicas and reported as mean $\pm$ standard deviation.

### 4 Supplementary Results

#### 4.1 Spectral Correlations and Regime Distributions

Figure S5 presents the complete, hierarchically clustered correlation matrix among all extracted spectral features, the Sequentiality Index, behavioural metrics, initialization scheme, and task complexity, which is defined as an ordinal variable (1–4) assigned a priori to each decision-making task, reflecting the presumed cognitive demand. A value of 1 corresponds to the simplest delayed binary decision; 2 adds context-dependent processing; 3 marks tasks requiring multiple distinct time intervals; and 4 denotes evidence-integration tasks that demand accumulation of information over time. This fixed mapping allows a straightforward exploration of how spectral and behavioural properties of trained networks scale with task difficulty. Figure S6 shows the distribution of key spectral quantities grouped by the dynamical regime (Oscillatory, Mixed, Ramping/Decaying).Together with the main figures, these results show that the maximum imaginary part of the recurrent weight eigenvalues has the largest point-biserial correlation with the oscillatory label among the descriptors examined, although no individual descriptor, including this one, remains significant after permutation testing and FDR correction (main text Fig. 9A). Initialization scheme reliably predicts the non-normality of the trained recurrent matrix (orthogonal initialization yields a near-zero Henrici index), consistent with orthogonal matrices being normal by construction, but its association with the final dynamical regime is comparatively weak (Table 2, main text). Jointly, however, the spectral descriptors are informative: a logistic classifier combining them discriminates oscillatory from other networks above a task-stratified permutation null (main text Fig. 9B), indicating that the final connectivity retains some regime-related information distributed across descriptors, without any single feature being individually predictive.

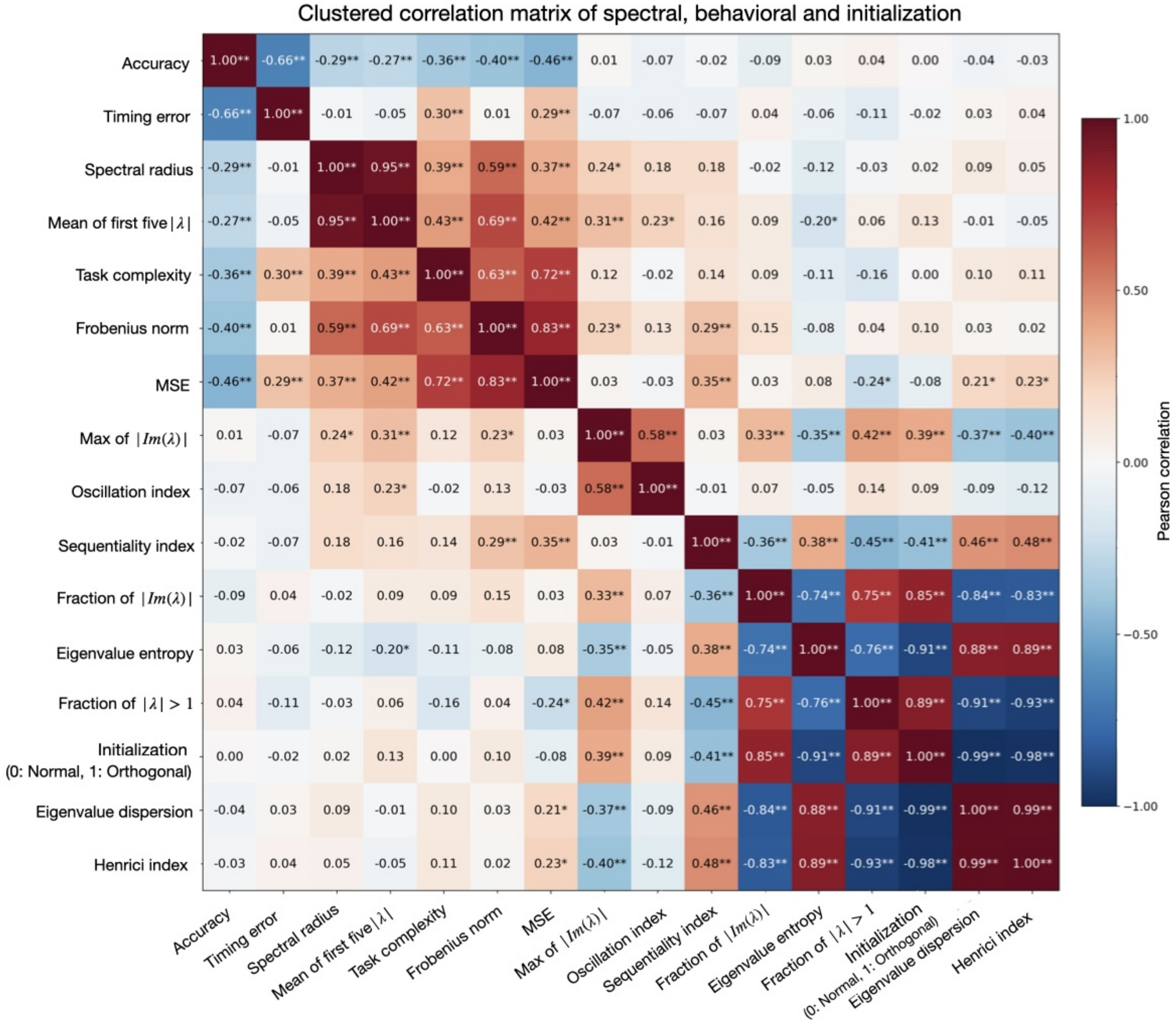


Figure S5: **Clustered correlation matrix of spectral, behavioural, and initialization variables.** Pearson correlation coefficients computed across all networks ($n = 100$). Rows and columns are reordered by hierarchical clustering to reveal blocks of strongly correlated quantities. Colour bar: Pearson correlation coefficient. Significance levels are indicated as $^{*}p < 0.05$, $^{**}p < 0.01$. The strong negative correlation between orthogonal initialization (Init $= 1$) and the Henrici index ($r = -0.98$) and the positive correlation between maximum imaginary part and the Oscillation Index ($r = 0.58$).

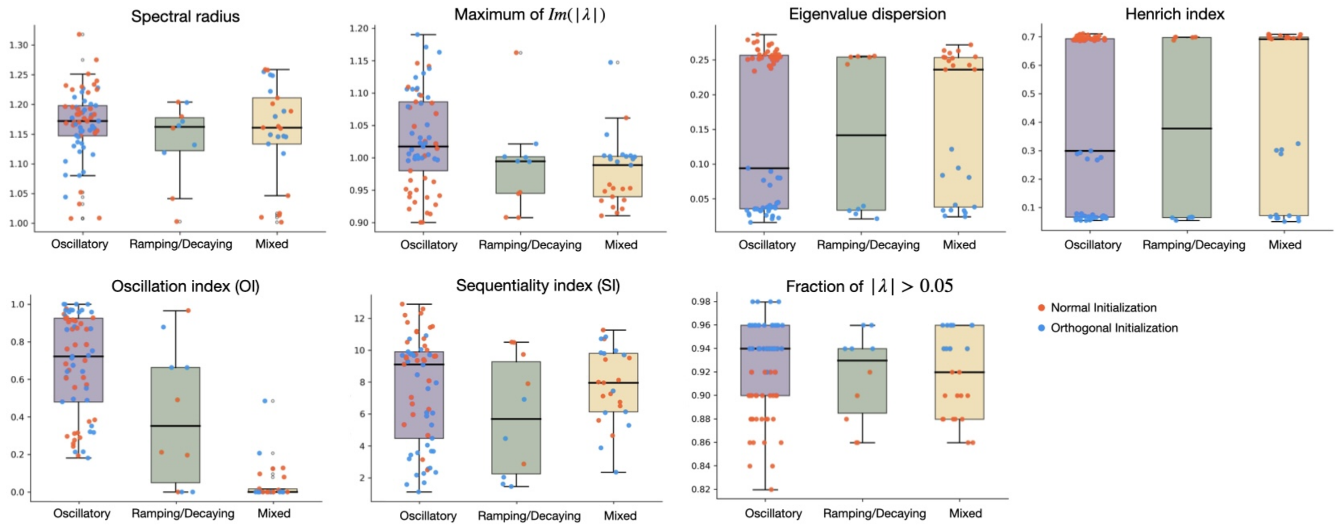


Figure S6: **Spectral features and Sequentiality Index grouped by dynamical regime.** Boxplots for each quantity (spectral radius, maximum imaginary part, eigenvalue dispersion, Henrici index, Oscillation Index, Sequentiality Index, and fraction of eigenvalues with $|\mathrm{Im}(\lambda)| > 0.05$) for networks classified as Oscillatory, Mixed, or Ramping/Decaying. Individual replicas are shown as dots, coloured by initialization scheme (orange: Normal, blue: Orthogonal). Box colors represent the three regimes (purple: Oscillatory, green: Ramping/Decaying, amber: Mixed). Oscillatory networks consistently exhibit higher maximum imaginary part and Oscillation Index, while Mixed networks span a wide range of Sequentiality Index values.

#### 4.2 Explained variance and Population geometry analysis.

To characterize condition-dependent population structure, we analyzed trial-averaged hidden-state activity using principal component analysis (PCA). For each condition and task stage (pre-stimulus, stimulus, intermediate, decision), we computed the leading three principal components of the time-by-neuron activity matrix, defining a low-dimensional subspace capturing dominant population variance. Subspace alignment between conditions was quantified using principal angles obtained from the singular value decomposition of the corresponding PCA bases. All analyses were performed across multiple independently trained network replicas, and results are reported as means and standard deviations across them. In addition, the explained variance ratio (EVR) of the leading components was computed from full-trial activity to assess the distribution of variance across population modes.

Across all tasks, population activity was well captured by a low-dimensional subspace, with the leading components accounting for the majority of variance. The alignment between condition-specific subspaces depended strongly on task structure and processing stage. In tasks where the stimulus directly specified the required response (e.g., sign or amplitude), subspace separation emerged during the stimulus epoch and persisted through the decision phase. In contrast, in tasks requiring temporal integration, subspaces remained aligned during the stimulus period and diverged only at later stages, near the time of response generation. These results indicate that the timing of subspace separation reflects whether task-relevant information is immediately available or must be accumulated over time.

##### 4.2.1 Delayed Decision-making Task

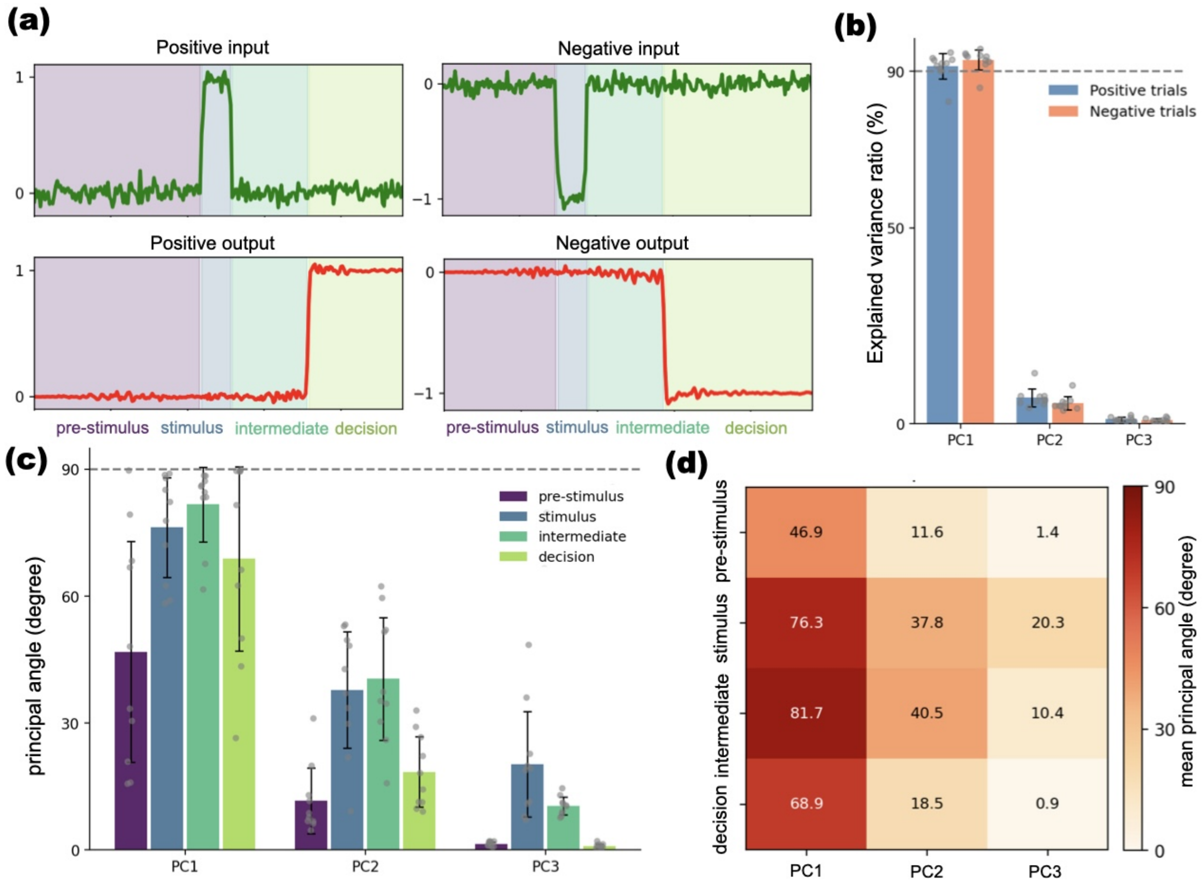


Figure S7: **Population geometry in the simple delayed decision-making task.** **(a)** Representative input (green) and network output (red) traces for positive and negative trials in the simple delayed decision-making task. Shaded regions indicate task phases (pre-stimulus, stimulus, intermediate, and decision). **(b)** Explained variance ratio (EVR) of the first three principal components computed from full-trial activity, shown separately for each condition. **(c)** Principal angles between the leading three-dimensional PCA subspaces of trial-averaged population activity for positive and negative conditions, computed across task-relevant stages: (pre-stimulus, stimulus, intermediate, decision). Bars denote means across network replicas; points indicate individual replicas. **(d)** Heatmap of mean principal angles as a function of epoch and principal component, summarizing subspace alignment across conditions.

| **Condition** | **PC1** | **PC2** | **PC3** |
|---|---|---|---|
| Positive | 91.3± 3.3% | 6.7± 2.3% | 1.2± 0.5% |
| Negative | 93.0± 2.6% | 5.2± 1.7% | 1.0± 0.4% |

Table S2: Explained Variance Ratio (full trial) for Simple Delayed Decision-making Task.

| Phase | PC1 | PC2 | PC3 |
|---|---|---|---|
| pre-stimulus | 46.9 ±26.1 | 11.6± 7.7 | 1.4± 0.5 |
| stimulus | 76.3± 11.8 | 37.8± 13.8 | 20.3± 12.4 |
| intermediate | 81.7±8.8 | 40.5±14.5 | 10.4± 2.1 |
| decision | 68.9±21.8 | 18.5±8.3 | 0.9± 0.5 |
| full trial | 3.8± 2.8 | 1.6± 0.5 | 0.1± 0.0 |

Table S3: Principal angles (degrees) between positive and negative subspaces for Simple Delayed Decision-making Task. Mean ± SD across replicas.

###### 4.2.2 Windowed Evidence Integration Task

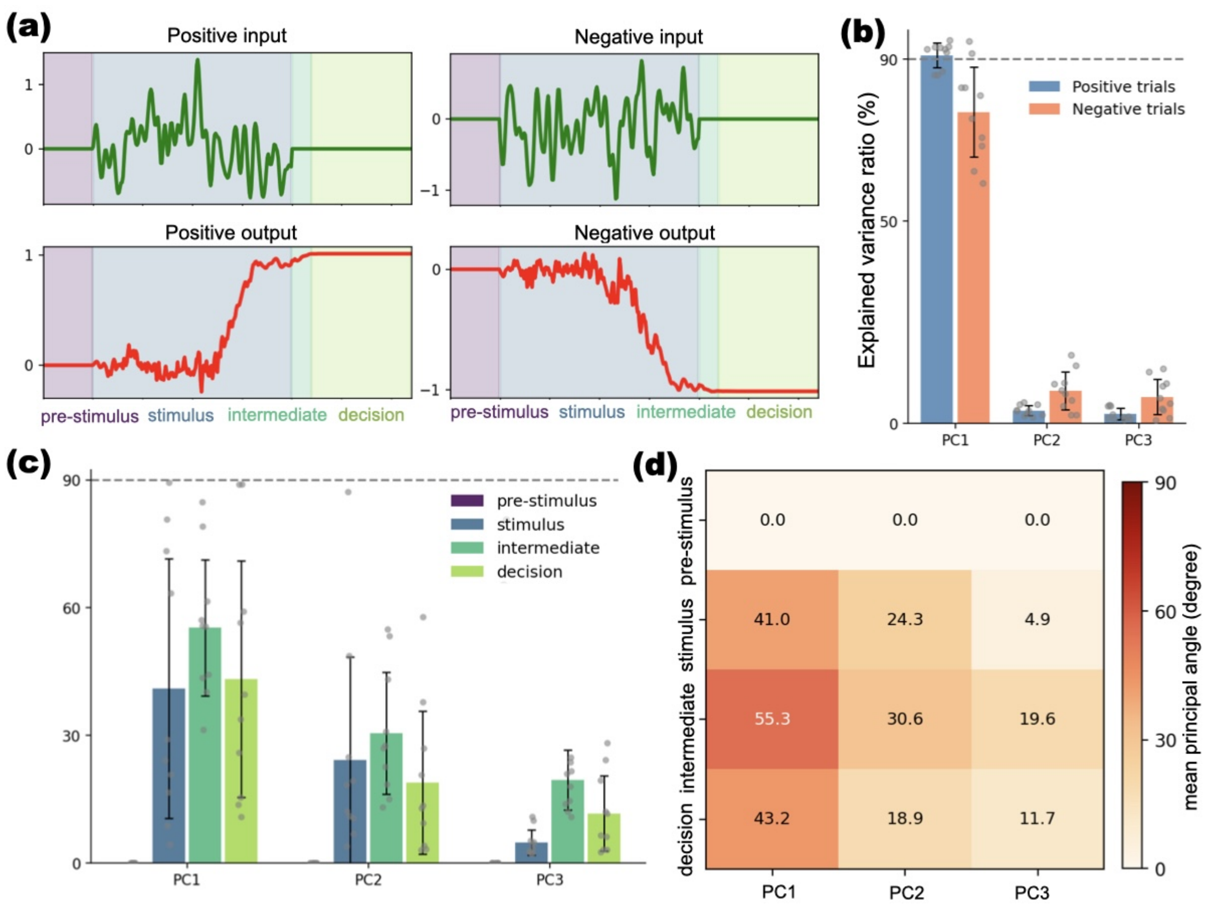


Figure S8: **Population geometry in the perceptual integration task (Integral DM). (a)** Representative input (green) and network output (red) traces for positive and negative trials in the integration DM task. Shaded regions indicate task phases (pre-stimulus, stimulus, intermediate, and decision). **(b)** Explained variance ratio (EVR) of the leading principal components computed from full-trial activity. **(c)** Principal angles between PCA subspaces associated with positive and negative accumulated evidence, computed across task stages. Subspace alignment is preserved during the stimulus (integration) period and diverges during intermediate and decision stages. Bars show means across replicas; points denote individual networks. **(d)** Heatmap of mean principal angles across epochs and components, highlighting the delayed emergence of subspace separation.

| Condition | PC1 | PC2 | PC3 |
|---|---|---|---|
| Positive | 91.0± 3.1% | 3.2± 1.2% | 2.4± 1.4% |
| Negative | 77.0± 11.1% | 8.1± 4.6% | 6.6± 4.3% |

Table S4: Explained Variance Ratio (full trial) for the Windowed Evidence Integration Task.

| **Phase** | **PC1** | **PC2** | **PC3** |
|---|---|---|---|
| pre-stimulus | 0.0± 0.0 | 0.0± 0.0 | 0.0± 0.0 |
| stimulus | 41.0± 30.5 | 24.3± 24.2 | 4.9± 3.0 |
| intermediate | 55.3±16.0 | 30.6± 14.3 | 19.6± 7.0 |
| decision | 43.2±29.6 | 18.9±17.5 | 11.7± 1.1 |

Table S5: Principal angles (degrees) between positive and negative subspaces for the Windowed Evidence Integration Task. Mean ± SD across replicas.

##### 4.2.3 Multi-interval amplitude-based DM

In the multi-interval amplitude task, the amplitude of the input pulse (A1 to A8) parametrically sets the duration of the delay period that the network must maintain before generating the response. A1 (lowest amplitude) engages the shortest delay of 25 ms after stimulus offset, A2 requires 50 ms, and the delay increases linearly by 25 ms per level up to A8 (highest amplitude) at 200 ms. Thus, larger pulse amplitudes demand progressively longer memory intervals, allowing the network's temporal processing to be examined across a range of delays

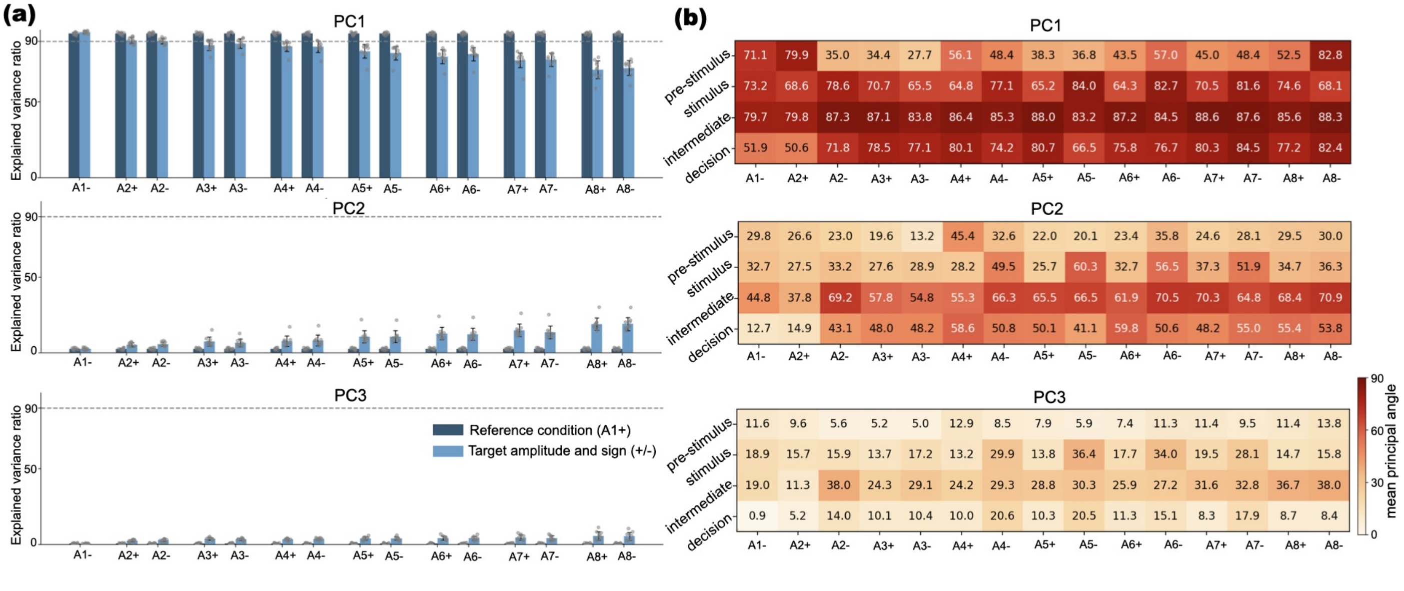

Figure S9: **Population geometry across amplitude-coded intervals.** **(a)** Explained variance ratio (EVR) of the leading principal components for each condition. **(b)** Principal angles between subspaces associated with different amplitude conditions summarized across each task stage. **(a)** and **(b)** correspond to the different PCA projections.

Table S6: Explained Variance Ratio (full trial) – Multi-amplitude task (Simple DM 8 times). Mean ± SD across replicas.

| **Positive amplitudes** | | | |
|---|---|---|---|
| **Condition** | **PC1** | **PC2** | **PC3** |
| A1+ | 95.2±0.7% | 2.6±0.6% | 0.8±0.3% |
| A2+ | 90.6±2.1% | 5.4±1.2% | 2.6±0.8% |
| A3+ | 87.4±3.5% | 7.5±3.0% | 3.9±0.8% |
| A4+ | 86.7±3.5% | 7.7±3.5% | 3.5±0.6% |
| A5+ | 83.4±4.5% | 10.7±4.0% | 4.1±1.1% |
| A6+ | 79.8±4.6% | 12.8±4.1% | 4.3±1.4% |
| A7+ | 77.5±4.8% | 15.0±4.1% | 4.7±1.8% |
| A8+ | 71.3±5.9% | 19.0±4.1% | 5.7±2.7% |

| **Negative amplitudes** | | | |
|---|---|---|---|
| **Condition** | **PC1** | **PC2** | **PC3** |
| A1- | 96.1±0.8% | 2.6±0.7% | 0.7±0.3% |
| A2- | 90.1±1.9% | 5.8±1.4% | 2.9±0.9% |
| A3- | 88.5±3.2% | 6.6±2.5% | 3.5±0.7% |
| A4- | 86.4±3.8% | 8.2±3.6% | 3.7±0.7% |
| A5- | 82.1±4.5% | 10.7±3.9% | 4.0±1.1% |
| A6- | 81.5±4.4% | 12.3±4.1% | 4.2±1.3% |
| A7- | 77.8±4.4% | 13.6±4.2% | 4.3±1.5% |
| A8- | 72.4±4.8% | 19.1±4.3% | 5.5±2.7% |

Table S7: Principal angles (degrees) – **PC1**. Reference condition A1+ (angles with itself are not applicable, shown as "–"). Mean ± SD.

| **Positive amplitudes (reference A1+)** | | | | | | | | |
|---|---|---|---|---|---|---|---|---|
| **Phase** | **A1+** | **A2+** | **A3+** | **A4+** | **A5+** | **A6+** | **A7+** | **A8+** |
| pre-stimulus | – | 79.9±10.1 | 34.4±28.4 | 56.1±33.2 | 38.3±25.8 | 43.5±29.5 | 45.0±23.5 | 52.5±22.9 |
| stimulus | – | 68.6±15.0 | 70.7±14.7 | 64.8±18.0 | 65.2±17.7 | 64.3±18.1 | 70.5±14.3 | 74.6±12.8 |
| intermediate | – | 79.8±11.7 | 87.1± 2.8 | 86.4± 3.6 | 88.0± 1.8 | 87.2± 2.5 | 88.6± 1.6 | 85.6± 7.1 |
| decision | – | 50.6±19.9 | 78.5±14.3 | 80.1±12.5 | 80.7±11.8 | 75.8±17.1 | 80.3±13.7 | 77.2±18.1 |

| **Negative amplitudes** | | | | | | | | |
|---|---|---|---|---|---|---|---|---|
| **Phase** | **A1-** | **A2-** | **A3-** | **A4-** | **A5-** | **A6-** | **A7-** | **A8-** |
| pre-stimulus | 71.1±14.9 | 35.0±31.0 | 27.7±26.1 | 48.4±33.8 | 36.8±26.9 | 57.0±31.6 | 48.4±27.5 | 82.8± 9.3 |
| stimulus | 73.2±13.7 | 78.6± 8.5 | 65.5±13.7 | 77.1±16.4 | 84.0±13.4 | 82.7± 7.2 | 81.6± 9.2 | 68.1±15.7 |
| intermediate | 79.7± 7.3 | 87.3± 1.8 | 83.8± 8.7 | 85.3± 8.6 | 83.2±12.7 | 84.5±12.1 | 87.6± 2.3 | 88.3± 1.6 |
| decision | 51.9±21.1 | 71.8±17.1 | 77.1±16.5 | 74.2±18.2 | 66.5±14.0 | 76.7±15.3 | 84.5± 3.1 | 82.4±13.3 |

Table S8: Principal angles (degrees) – **PC2**. Reference condition A1+ (angles with itself are not applicable, shown as "–"). Mean ± SD.

| Positive amplitudes (reference A1+) | | | | | | | | |
|---|---|---|---|---|---|---|---|---|
| **Phase** | **A1+** | **A2+** | **A3+** | **A4+** | **A5+** | **A6+** | **A7+** | **A8+** |
| pre-stimulus | – | 26.6±34.9 | 19.6±30.6 | 45.4±40.9 | 22.0±31.4 | 23.4±30.7 | 24.6±30.5 | 29.5±31.9 |
| stimulus | – | 27.5±29.8 | 27.6±27.9 | 28.2±23.2 | 25.7±22.1 | 32.7±22.5 | 37.3±24.5 | 34.7±25.4 |
| intermediate | – | 37.8±26.1 | 57.8±21.3 | 55.3±25.3 | 65.5±19.1 | 61.9±21.8 | 70.3±17.5 | 68.4±21.5 |
| decision | – | 14.9± 5.1 | 48.0±17.1 | 58.6±21.7 | 50.1±18.9 | 59.8±17.3 | 48.2±19.3 | 55.4±21.6 |

| Negative amplitudes | | | | | | | | |
|---|---|---|---|---|---|---|---|---|
| **Phase** | **A1-** | **A2-** | **A3-** | **A4-** | **A5-** | **A6-** | **A7-** | **A8-** |
| pre-stimulus | 29.8±28.8 | 23.0±32.9 | 13.2±18.3 | 32.6±34.0 | 20.1±32.5 | 35.8±33.0 | 28.1±31.9 | 30.0±32.5 |
| stimulus | 32.7±29.7 | 33.2±26.9 | 28.9±11.7 | 49.5±24.5 | 60.3±25.2 | 56.5±21.3 | 51.9±23.1 | 36.3±25.2 |
| intermediate | 44.8±31.0 | 69.2±17.9 | 54.8±22.4 | 66.3±21.0 | 66.5±24.1 | 70.5±22.5 | 64.8±19.7 | 70.9±16.2 |
| decision | 12.7±13.9 | 43.1±14.5 | 48.2±19.7 | 50.8±16.7 | 41.1±10.1 | 50.6±17.4 | 55.0±20.0 | 53.8±22.0 |

Table S9: Principal angles (degrees) – **PC3**. Reference condition A1+ (angles with itself are not applicable, shown as "–"). Mean ± SD.

| Positive amplitudes (reference A1+) | | | | | | | | |
|---|---|---|---|---|---|---|---|---|
| **Phase** | **A1+** | **A2+** | **A3+** | **A4+** | **A5+** | **A6+** | **A7+** | **A8+** |
| pre-stimulus | – | 9.6±17.0 | 5.2± 4.2 | 12.9±10.0 | 7.9± 6.8 | 7.4± 5.7 | 11.4±11.6 | 11.4±11.2 |
| stimulus | – | 15.7±21.9 | 13.7±13.6 | 13.2±12.0 | 13.8±10.2 | 17.7±17.3 | 19.5±18.4 | 14.7± 9.9 |
| intermediate | – | 11.3± 7.4 | 24.3±19.8 | 24.2±12.9 | 28.8± 9.8 | 25.9±12.0 | 31.6±13.8 | 36.7±12.8 |
| decision | – | 5.2± 2.6 | 10.1± 4.3 | 10.0± 3.4 | 10.3± 3.2 | 11.3± 3.6 | 8.3± 2.4 | 8.7± 2.1 |

| Negative amplitudes | | | | | | | | |
|---|---|---|---|---|---|---|---|---|
| **Phase** | **A1-** | **A2-** | **A3-** | **A4-** | **A5-** | **A6-** | **A7-** | **A8-** |
| pre-stimulus | 11.6±18.0 | 5.6± 4.1 | 5.0± 2.2 | 8.5± 7.7 | 5.9± 6.1 | 11.3± 7.1 | 9.5± 8.7 | 13.8±21.9 |
| stimulus | 18.9±18.5 | 15.9±13.6 | 17.2±10.3 | 29.9±20.5 | 36.4±15.4 | 34.0±13.7 | 28.1±14.5 | 15.8±12.4 |
| intermediate | 19.0±21.5 | 38.0±11.8 | 29.1±13.1 | 29.3±12.6 | 30.3±15.1 | 27.2±10.3 | 32.8±11.9 | 38.0±17.5 |
| decision | 0.9± 0.5 | 14.0± 2.5 | 10.4± 4.8 | 20.6± 8.6 | 20.5± 7.6 | 15.1± 7.3 | 17.9± 7.1 | 8.4± 3.1 |

##### 4.2.4 Multi-interval Distance-based Decision Making

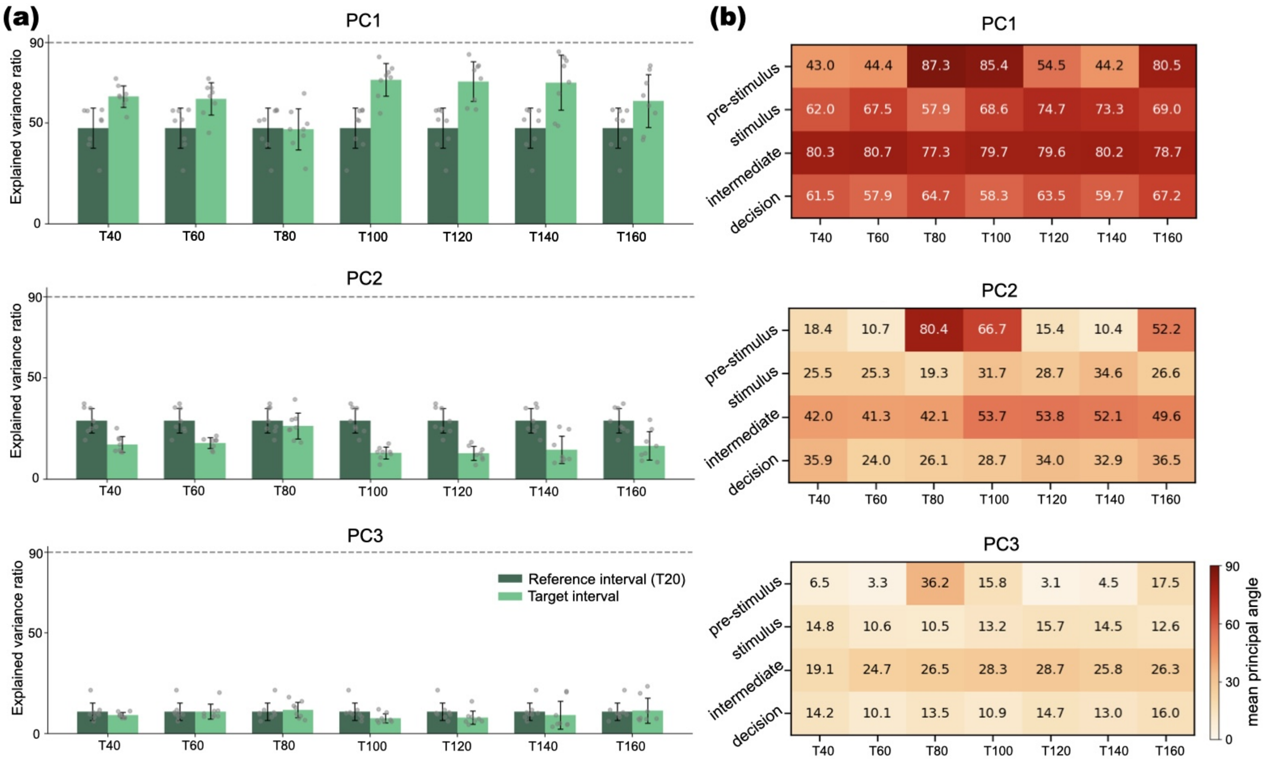


Figure S10: **Population geometry across interval-coded timing.** **(a)** Explained variance ratio (EVR) of the leading principal components for each condition. **(b)** Principal angles between subspaces associated with different interval conditions summarized across each task stage. **(a)** and **(b)** correspond to the different PCA projections.

| **Condition** | **PC1** | **PC2** | **PC3** |
|---|---|---|---|
| $T_{20}$ | 45.8± 9.1% | 25.2± 4.5% | 14.8± 4.0% |
| $T_{40}$ | 48.0± 9.2% | 23.9± 3.8% | 12.7± 4.1% |
| $T_{60}$ | 71.5± 5.7% | 12.3± 2.4% | 8.9± 2.5% |
| $T_{80}$ | 53.5±10.8% | 23.3± 7.4% | 12.5± 3.5% |
| $T_{100}$ | 80.7± 4.4% | 8.9± 2.4% | 5.4± 1.3% |
| $T_{120}$ | 70.9±11.6% | 14.0± 4.6% | 9.0± 5.6% |
| $T_{140}$ | 58.0±13.6% | 20.4±10.1% | 12.0± 4.6% |
| $T_{160}$ | 65.2±14.3% | 15.3± 6.9% | 10.3± 6.2% |

Table S10: Explained Variance Ratio (full trial) – Simple DM 8 time encoded task. Mean ± SD across replicas.

Table S11: Principal angles (degrees) – **PC1**. Reference T20 vs. each interval. Mean ± SD.

| **Phase** | $T_{40}$ | $T_{60}$ | $T_{80}$ | $T_{100}$ | $T_{120}$ | $T_{140}$ | $T_{160}$ |
|---|---|---|---|---|---|---|---|
| pre-stimulus | 43.0±27.7 | 44.4±25.8 | 87.3± 1.9 | 85.4± 6.5 | 54.5±26.6 | 44.2±25.6 | 80.5± 9.0 |
| stimulus | 62.0±17.1 | 67.5±18.6 | 57.9±22.9 | 68.6±17.6 | 74.7±12.6 | 73.3±17.8 | 69.0±18.3 |
| intermediate | 80.3±12.1 | 80.7±15.2 | 77.3± 7.5 | 79.7± 8.2 | 79.6± 9.8 | 80.2±10.8 | 78.7± 9.0 |
| decision | 61.5±21.5 | 57.9±19.6 | 64.7±12.4 | 58.3±21.1 | 64.5±20.9 | 59.8±22.7 | 67.4±22.6 |

Table S12: Principal angles (degrees) – **PC2**. Reference T20 vs. each interval. Mean ± SD.

| **Phase** | $T_{40}$ | $T_{60}$ | $T_{80}$ | $T_{100}$ | $T_{120}$ | $T_{140}$ | $T_{160}$ |
|---|---|---|---|---|---|---|---|
| pre-stimulus | 18.4±14.9 | 10.7±12.0 | 80.4± 2.9 | 66.7± 8.3 | 15.4±15.4 | 10.4±10.8 | 52.2±14.5 |
| stimulus | 25.5±10.6 | 25.3± 6.0 | 19.3± 5.1 | 31.7±13.3 | 28.7± 9.3 | 34.6±11.3 | 26.6± 9.4 |
| intermediate | 42.0±14.9 | 41.3± 7.0 | 42.1±11.3 | 53.7±12.4 | 53.8±19.5 | 52.1±20.3 | 49.6±16.7 |
| decision | 35.9±21.9 | 24.0±11.2 | 26.1±11.8 | 28.7±14.5 | 34.0±26.5 | 32.9±20.8 | 36.4±24.5 |

Table S13: Principal angles (degrees) – **PC3**. Reference T20 vs. each interval. Mean ± SD.

| **Phase** | $T_{40}$ | $T_{60}$ | $T_{80}$ | $T_{100}$ | $T_{120}$ | $T_{140}$ | $T_{160}$ |
|---|---|---|---|---|---|---|---|
| pre-stimulus | 6.5± 6.3 | 3.3± 1.2 | 36.2± 7.0 | 15.8± 4.6 | 3.1± 0.9 | 4.5± 2.8 | 17.5±11.5 |
| stimulus | 14.8± 6.7 | 10.6± 3.4 | 10.5± 4.0 | 13.2± 4.4 | 15.7± 6.6 | 14.5± 6.5 | 12.6± 3.3 |
| intermediate | 19.1± 6.9 | 24.7± 9.0 | 26.5± 8.0 | 28.3± 9.5 | 28.7±11.4 | 25.8± 7.2 | 26.3± 8.1 |
| decision | 14.2± 7.3 | 10.1± 4.9 | 13.5± 4.4 | 10.9± 4.1 | 14.7±11.7 | 13.0± 3.9 | 16.0±11.7 |

#### 4.3 Task generalization

Networks trained on eight discrete stimulus amplitudes (1–8) achieved perfect accuracy ($1.00 \pm 0.00$) for all interpolation values within that range, with relative timing errors of $-8.96 \pm 12.98\%$ (Normal init.) and $-4.20 \pm 6.51\%$ (Orthogonal init.), demonstrating that they faithfully learned the amplitude-to-delay mapping for the trained parameter region. Note that relative timing error was defined as $\frac{t_{\text{pred}} - t_{\text{target}}}{t_{\text{target}}} \times 100$, where $t_{\text{target}}$ is the time at which target output deviates from zero and $t_{\text{pred}}$ is the first time step at which the network output crosses a fixed threshold $\theta = 0.3$ in the same direction. Extrapolation below the training range (amplitudes $< 1$) produced a sharp drop in accuracy ($0.74 \pm 0.28$ and $0.53 \pm 0.23$ for Normal and Orthogonal, respectively) and a large increase in relative timing error (up to 41.75% for Normal), indicating that the networks cannot sustain the required longer delays for very weak stimuli. Extrapolation above the training range (amplitudes $> 8$) maintained relatively high accuracy ($0.92 \pm 0.14$ and $0.84 \pm 0.29$) but introduced a systematic negative timing bias (around $-18\%$), meaning the networks anticipated the response onset relative to the true delay. This dissociation is clearly visible in the example trace figures, where the gray target-delay arrows and the colored predicted-delay arrows diverge progressively outside the training region.

In contrast, in the windowed evidence-integration task, networks trained with a single 200-step integration window exhibited robust extrapolation across a wide range of window lengths (50–430 steps). Accuracy remained high (0.91–0.96), and relative onset errors were small (below 3% in absolute value), indicating that the learned integration strategy is largely scale-invariant. This result is consistent with the expectation that a network capable of computing a simple integral can generalize to any window size, and it is reflected both in the summary table and in the timing-error curves, which stay flat and close to zero throughout the entire extrapolation range. The example traces for this task confirm that the network's response onset shifts appropriately with window length, closely matching the target timing.

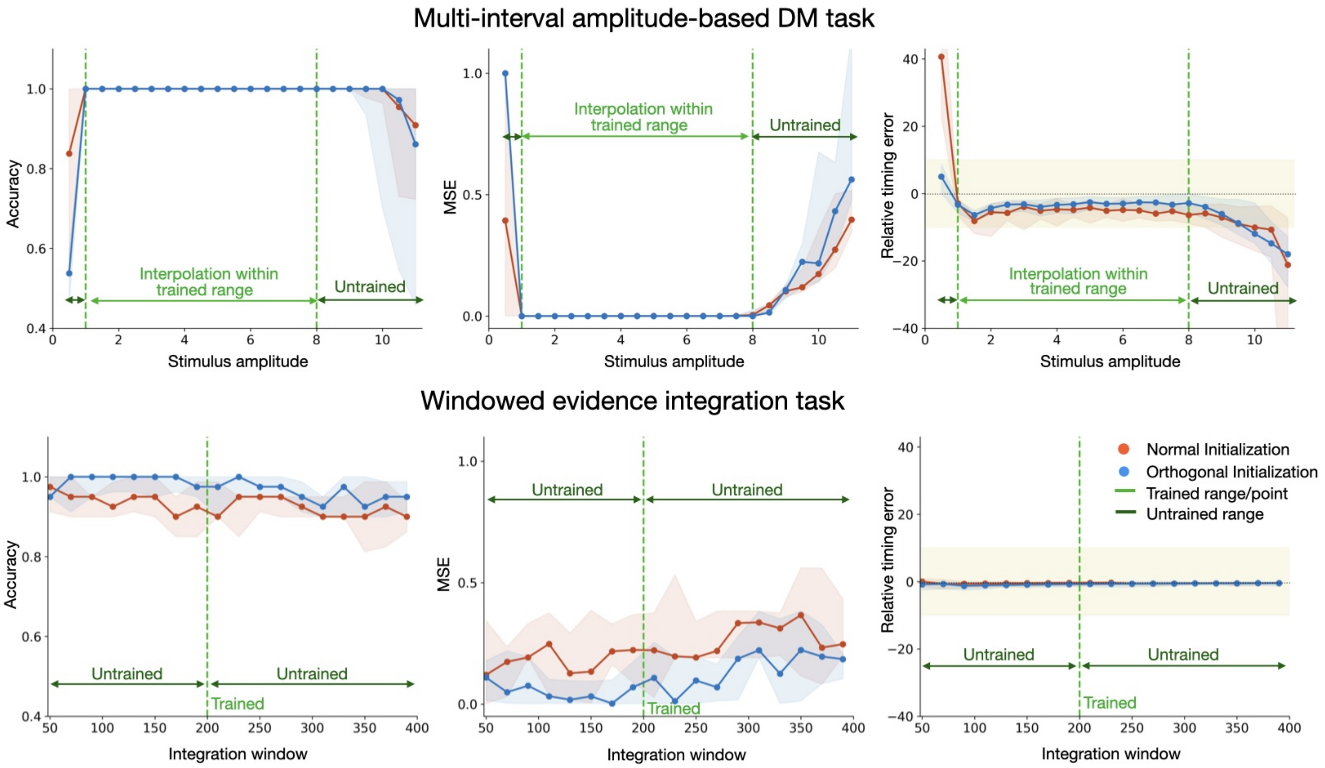


Figure S11: Accuracy (left), mean squared error (middle), and relative onset timing error (right) for the two tasks: multi-interval amplitude-based (top row) and windowed evidence integration (bottom row). In each panel, the training range (or single trained window for the integration task) is marked by dashed green vertical lines with horizontal arrows labeling the trained and untrained regions. Solid lines and shaded error bands represent the median and interquartile range (25th - 75th percentile) across network replicas, with orange for Normal and blue for Orthogonal initializations. In the relative timing error panels, the yellow shaded region spans 10%, and the dashed horizontal line indicates zero error. The amplitude-based task shows excellent performance within the trained range but a clear degradation outside it, both in accuracy and timing error. The integration task generalizes well across all tested window lengths, maintaining high accuracy and small timing errors even if every tested window length other than window value (200) falls in the untrained region.

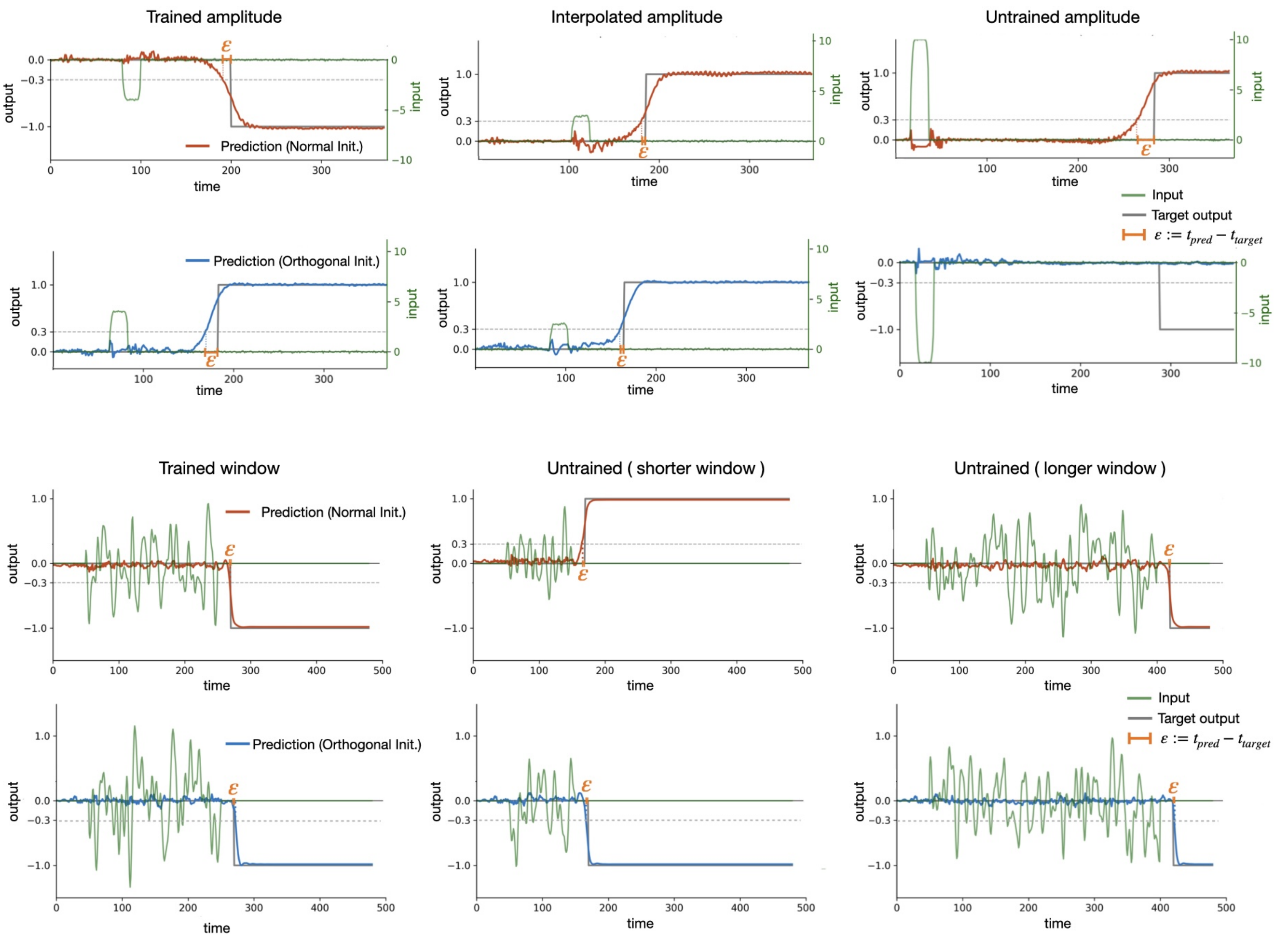


Figure S12: Example network output traces with stimulus input and response onset error for the two tasks: multi-interval amplitude-based (top two rows) and windowed evidence integration (bottom two rows). Each panel shows a single trial for one network replica (odd rows: Normal initialization; even rows: Orthogonal initialization). For each task, columns correspond to three representative parameter values: a trained value (left), an interpolated or shorter untrained value (middle), and an extrapolated or longer untrained value (right). The green line depicts the input stimulus, the gray line the target output, and the colored line the network's predicted output (orange: Normal, blue: Orthogonal). For the amplitude-based task, input is plotted on a separate right-hand axis (scaled to the stimulus amplitude); for the integration task, input and output share the same axis. The dashed horizontal lines mark the $\pm 0.3$ onset-detection threshold used to determine response onset. The onset timing error, $\varepsilon := t_{\text{pred}} - t_{\text{target}}$, is marked at the point where the network output first crosses this threshold. For the trained and interpolated/shorter parameter values, the predicted onset closely tracks the target in both tasks; for the extrapolated amplitude (top-right, bottom-right), the predicted onset deviates noticeably, illustrating the breakdown of the learned temporal mapping outside the trained range, whereas the integration task's predicted onset remains accurate even for the longer untrained window (bottom-right), reflecting the scale-invariance of the learned integration strategy.
